# Loop competition and extrusion model predicts CTCF interaction specificity

**DOI:** 10.1101/2020.07.02.185389

**Authors:** Wang Xi, Michael A. Beer

## Abstract

Three-dimensional chromatin looping interactions play an important role in constraining enhancer-promoter interactions and mediating transcriptional gene regulation. CTCF is thought to play a critical role in the formation of these loops, but the specificity of which CTCF binding events form loops and which do not is difficult to predict. Loops often have convergent CTCF binding site motif orientation, but this constraint alone is only weakly predictive of genome-wide interaction data. Here we present an easily interpretable and simple mathematical model of CTCF mediated loop formation which is consistent with Cohesin extrusion and can predict ChIA-PET CTCF looping interaction measurements with high accuracy. Competition between overlapping loops is a critical determinant of loop specificity. We show that this model is consistent with observed chromatin interaction frequency changes induced by CTCF binding site deletion, inversion, and mutation, and is also consistent with observed constraints on validated enhancer-promoter interactions.

## Introduction

High order chromatin structure affects various biological processes within the nucleus, ranging from gene regulation to DNA repair. The structural basis of interphase chromatin has been extensively studied by various Chromatin Conformation Capture^1–4^ techniques, and has revealed functional units including chromosome compartments^1^, topologically associated domains (TADs)^5^ and loops^6^. Chromosomal compartments, which exhibit a checkerboard pattern on a Hi-C map, correspond to active or inactive chromatin across several megabases^1^. On the other hand, TADs and sub-TAD loops represent enriched chromatin interactions that appear at a scale of hundreds of kilobases or below^5,6^. These smaller loops shape local chromatin structure, and their disruption has been reported to lead to dramatic dysregulation of nearby gene expression^7,8^. The most prominent feature of TADs and loops is that their boundaries are usually marked by CTCF and Cohesin binding^5,6^. CTCF was initially thought to work mainly as an insulator of active chromatin marks, but since has been recognized to play a major role in chromatin organization, whereby pairs of CTCFs bind and serve as loop anchors to constrain interactions between distant regulatory elements^9,10^ (Fig. 1a). It has been suggested that CTCF and Cohesin mediate TAD and loop formation through a loop extrusion mechanism, where Cohesin translocation generates a nascent chromatin loop until blocked by CTCF^11,12^ (Fig. 1b). Polymer simulations of a loop extrusion model successfully reconstructed TAD like structures, and predicted the impact of CTCF or Cohesin degradation on TAD strength^11,12^. Moreover, multiple experiments have validated *in vitro* that Cohesin is capable of moving through nucleosomal DNA^13^ and generating a growing DNA loop progressively as it moves^14,15^.

**Figure 1.**
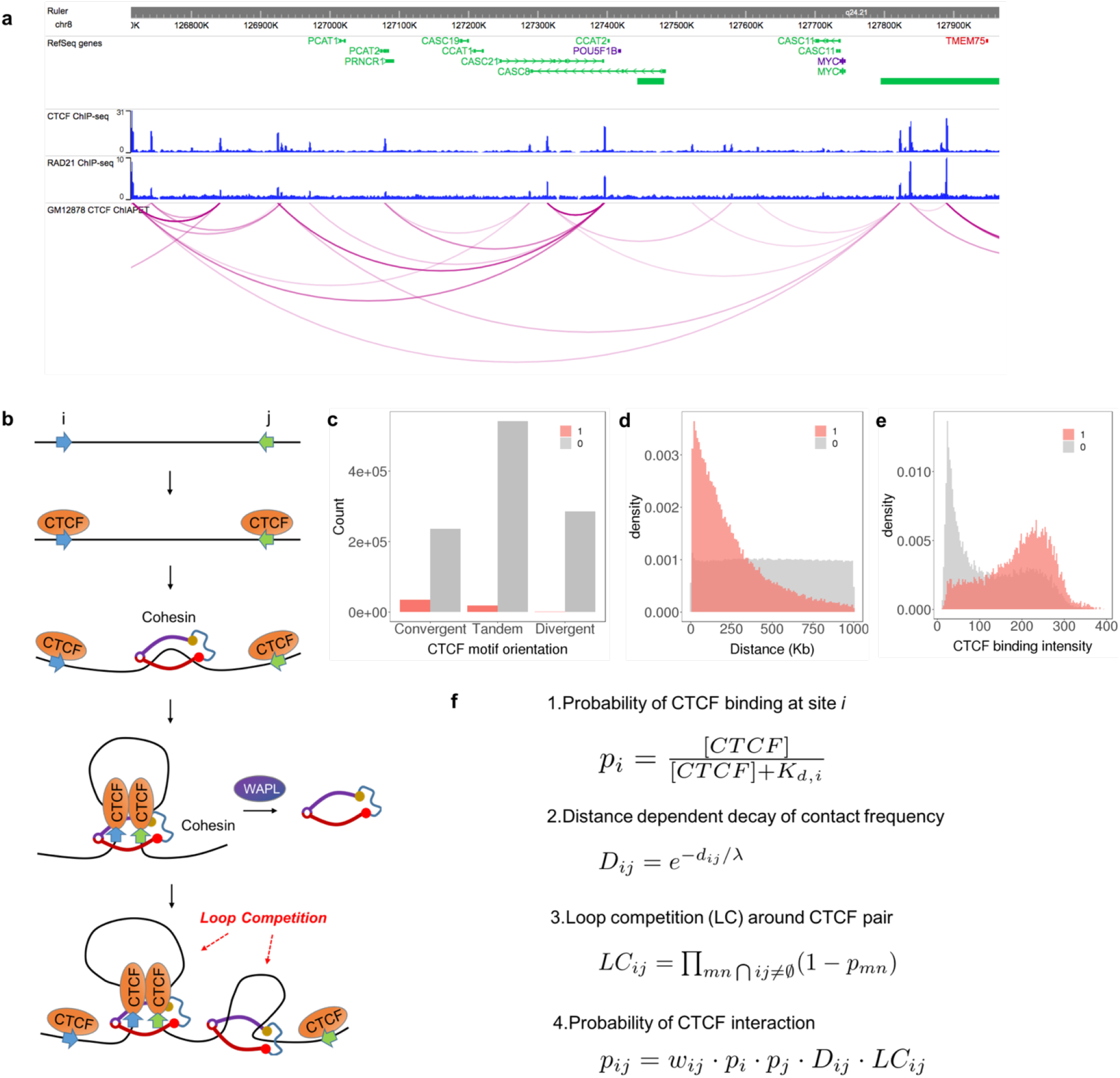
Mathematical formulation of a loop competition and extrusion model. (a) We use CTCF ChIA-PET to train our model: the contact profile of the Myc locus in GM12878 is shown. (b) Loop extrusion model: Cohesin is loaded between (typically convergent) CTCF pairs, and the loop forms progressively as Cohesin translocates along the chromatin fiber. The extrusion process stops when Cohesin is stalled by CTCF. WAPL unloads Cohesin from chromatin. An existing loop could block movement of another Cohesin protein, leading to loop competition. (c) While measured loops prefer convergent CTCF pairs, other orientations also interact with significant frequencies and many neighboring (<1Mb) convergent CTCF motifs do not form loops: shown are interacting pair counts (1, red), and non-interacting pair counts (0, grey). Here interacting and non-interacting loops are defined by ChIA-PET interaction data. (d) Distance/ distribution for interacting and non-interacting CTCF pairs. (e) CTCF binding intensity distribution for interaction and non-interaction CTCF pairs. (f) Mathematical model of loop interaction probability formed by this extrusion process.

There are ~50,000 CTCF binding sites in normal mammalian cells, which corresponds to over 1 million possible CTCF pairs lying within 1Mb of each other. However, only about 2~5% of these are identified to be interacting by direct Hi-C or ChIA-PET measurements^6,16^ (Fig. 1c). This raises the important question about the difference between interacting and non-interacting CTCF pairs. Although it has been observed that CTCF motif orientation in loop anchors tends to be convergent^6,17^, the vast majority of convergent CTCF motif pairs are not interacting with each other, therefore, a more comprehensive model of how CTCF interaction specificity is regulated remains to be elucidated. Several experiments have investigated the determinants of loop formation, such as, binding of CTCF or Cohesin^17–20^(Supplementary Fig.1), but could not explain why only a subset of available CTCF binding pairs are interacting in each cell type. While CTCF and Cohesin have been shown to play a role in determining 3D chromatin interactions overall, the process of loop extrusion has not yet been directly validated to be the molecular mechanism underlying CTCF looping interactions. Previous physical modeling of nuclear organization has focused more on general principles associated with the formation of TADs and loops but has not explored the variation in the strength of such features observed across different loci in real datasets^11,12^. Additionally, polymer physics based models treating the chromatin fiber as a connected chain of interacting units have shown encouraging global correspondence with measured contract frequencies, but their ability to predict individual CTCF loops has not been systematically evaluated^21–25^. In contrast, one machine learning model, Lollipop, utilized a large set of genomic and epigenomic features to predict specific CTCF interactions with high accuracy^26^. This model motivated our approach, and provides some insight into this problem, but did not fully reveal how these features play a role in the process of loop formation. Moreover, inspection of the Lollipop model shows that many of the 77 features used have substantial redundancy, making it hard to distinguish causal mechanisms, and implying that there may be simpler rules driving the specificity of CTCF interactions.

Here, we propose that CTCF interaction specificity can be predicted by a simple model based on loop extrusion. The success of this model gives indirect support for loop extrusion as an important mechanism regulating CTCF interaction specificity. We build a quantitative model to describe CTCF-mediated loop formation with only four features, CTCF binding intensity (BI), CTCF motif orientation, distance between CTCF binding events, and loop competition (LC) (Fig. 1c-1e). We show that this model can predict both ChIA-PET and Micro-C annotated CTCF loops with high accuracy. Our model includes an explicit contribution from the competition between overlapping loops, which is crucial for accurate prediction of loop formation. Our model of loop competition also provides a simple mechanism by which genetic variation in CTCF binding sites directly contributes to observed differences in chromatin contact frequency. We show that our model is also predictive of cell-type specific CTCF loops. We further validate this model by predicting published CRISPRi perturbations of loop anchor binding sites, and by the predicted CTCF loops’ ability to constrain enhancer promoter interactions. We expect that the insights derived from this model may also shed light onto the related important problem of enhancer-promoter interaction prediction, and the mechanisms by which the specificity of enhancer-promoter interactions are regulated.

## Results

### Quantitative model of loop formation by extrusion

In this loop extrusion model, the key components are CTCF, Cohesin and other loop-extruding factors^11,12^ (Fig. 1b). Our model relies on the assumption that looping interactions found in the CTCF ChIA-PET experiments are due to the blocking and localization of Cohesin at CTCF binding sites. The formation of a CTCF-mediated loop in mammalian cells begins when the ring-shaped Cohesin is loaded onto the DNA chromatin fiber. Through the motor activity of Cohesin and other co-factors like NIPBL, Cohesin translocate along the chromatin fiber in an ATP-dependent manner, which pushes and progressively enlarges the DNA loop. This process proceeds until Cohesin dissociates from DNA or comes into contact with a DNA bound CTCF protein on each strand of the loop, which acts as a barrier that prevents further translocation. That CTCF acts as a blockade to Cohesin and acts as primary determinant of genomic locations enriched in Cohesin in the genome is supported by gkm-SVM sequence analysis showing that the CTCF binding site alone is able to explain genomic binding of SMC3, a Cohesin subunit.^27^ The most stable loop configuration is thus a Cohesin bound DNA loop with a CTCF bound at each base of the loop, and there is a notable preference for these CTCF binding sites to be in a convergent orientation.

We built a simple model which predicts the probability of formation for all possible loops by quantitatively combining the contribution of each step in this process. (Fig. 1f). First, the probability of CTCF binding at each genomic binding site is described by the chemical equilibrium:

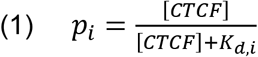

where [CTCF] is the concentration of CTCF to be inferred, *K_d,i_* is the local dissociation constant at site *i.*^28^ We will use the local ChIP-seq signal, *x,* to determine *K_d,i_*. To normalize, we let *x* be the local CTCF binding intensity signal divided by its genome average. Since *K_d,i_* is the dissociation constant, *x* is inversely proportional to *K_d,i._*, but with some unknown scaling factor. Since [CTCF] is also a constant, we can combine [CTCF] and the ChIP-seq signal scaling factor to write *x*=*a*∙[CTCF]/*K_d,i_* or 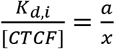, so the probability of binding, Eq. (1), can be simply written as: 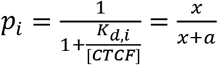. The dimensionless parameter *a* can be thought of as an estimate of the average 〈*K_d,i_*〉/[CTCF] over all the CTCF binding sites, and turns the local ChIP-seq signal intensity into a probability of occupancy. We will learn the best value of the parameter *a* from the ChIA-PET data. These binding probabilities contribute independently to a loop forming between CTCF site *i* and CTCF site *j.* In addition to the binding probability at each potential loop anchor site, we account for the contribution of CTCF motif orientation on loop stability with a scalar, *w_ij_*, and this term takes three different values, 1, 1/*w*, and 1/*w*^2^, for convergent, tandem or divergent CTCF motifs^17,19^. This simple one parameter orientation effect model is consistent with a more general treatment described in Methods and Supplementary Fig. 4a. The extrusion process adds an additional term which reflects the probability that Cohesin does not stochastically dissociate from the DNA fiber while translocating along it. A constant dissociation rate leads to an exponential decay term of the form:

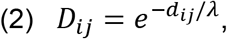

where *d_ij_* is the distance between CTCF sites *i* and *j*. For example, if the probability of not falling off while translocating 1 bp is *α*, the probability of not falling off after translocating *n* bp is *α^n^*, and in terms of distance *e*^−1/*λ*^ = *α*. This term leads to decreased loop interaction frequency when the distance between two CTCF bound regions gets larger. The parameter *λ* can also be interpreted as the processivity of Cohesin, or equivalently, the average CTCF loop length, which has been estimated to be about 300kb^11^.

The final notable component of our loop competition and extrusion model is the effect of loop competition. The mechanism of loop extrusion implies that one Cohesin bound loop could block additional Cohesin procession. This blocking prevents all CTCF pairs that overlap with a formed loop from interacting, since other Cohesins would have difficulty passing through, no matter where they load^11,12^. We will consider both a complete blocking model, where the presence of one loop excludes the formation of all overlapping loops, and an incomplete blocking model, where Cohesin can process through existing bound Cohesins with some probability. Allowing some pass-through is motivated by emerging evidence from a structurally similar loop extrusion factor, Condensin^29^, which we will also discuss in the context of a WAPL knockout. In the complete blocking model, the formation of one loop excludes all other overlapping loops, so the contribution of loop competition is:

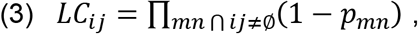

where *p_mn_* is the probability of loop formation between CTCF site *m* and *n*, as defined in Eq. 5 below. Specifically, *LC_ij_* is an additional contribution to *p_ij_* that reflects the constraint that an overlapping loop between two CTCF sites *m* and *n* is not formed. In this sense, the complete model with loop competition (Eq. 5, below) should be solved iteratively. But in Methods, we show that the full iterative solution of Eq. 3 is consistent with a simpler model, which just requires that all CTCF sites internal to the loop *ij* are unoccupied, using *p_m_* from Eq. 1 for the probability of occupancy of site *m*. This approximate loop competition model can thus be written:

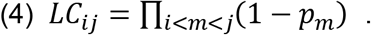

In practice this approximate loop competition term reflects the fact that strong sites inside a loop can contribute to internal loop formation and outcompete the formation of the loop *ij.* We assume that the probability of Cohesin loading is constant along genome, for the moment ignoring any non-uniformity or nuclear compartmentation. Thus in our complete model, the probability of a loop forming between CTCF binding sites *i* and *j* is given by:

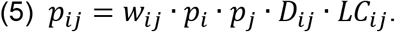

### Parameter determination for loop extrusion probabilistic model

We used publicly available CTCF ChIA-PET data^16^ in GM12878 and HeLa cells to determine the values of the parameters *a* = 〈*K_d,i_*〉/[CTCF], *w* and λ in our model. Long read ChIA-PET data was processed with ChIA-PET2 software under standard protocols to identify significant loops^30^. The high resolution and quality of this ChIA-PET data makes it suitable for predicting CTCF-mediated loops and training our model. First, the average anchor length of ChIA-PET loop is around 1kb, which is close to the size of open chromatin region around single CTCF binding site (Supplementary Fig. 2a). Second, comparison of CTCF ChIP-seq peaks with overlapping ChIA-PET anchors shows that they are relatively centered around each other (Supplementary Fig. 2b). We will use the CTCF ChIP-seq signal at each site as *K_d,i_* to infer the local CTCF binding probability. CTCF motif annotation is performed with STORM^31^.

We determined the optimal value of the model parameters by fitting the loop extrusion model to CTCF ChIA-PET data (Fig. 2a-2d, see Methods), by comparing measurements of actual loop formation to the probability of loop formation predicted by our model (AUPRC), using GM12878 and HeLa. The low dimensionality of our model makes overfitting highly unlikely, and training these three parameters on the full dataset or 5-fold cross validation both yield the same optimal values (Supplementary Fig. 3d-f). We did a comprehensive grid search in (〈*K_d,i_*〉/[CTCF], *w*, and λ) in GM12878 (Fig. 2d), and found that the *w* value of best agreement with data is 3.0, which implies that a convergent CTCF pair is three times more likely to interact than a tandem CTCF pair with equivalent CTCF binding probability and distance, and nine times more likely than a divergent pair. The optimal value of *a* = 〈*K_d,i_*〉/[CTCF] is 8.5. *K_d_* in vitro for CTCF binding to the H19/Igf2 CTCF binding site has been measured to be 370nM^32^ and nuclear [CTCF] is around 144nM^33–35^. This leads to *K_d_*/[CTCF] = 2.6. While our estimate of this parameter a=〈*K_d,i_*〉/[CTCF]=8.5 is near this value, it is not unreasonable to expect that the global average of *K_d_* at binding sites on chromatin in vivo will be somewhat higher than that measured on naked DNA in vitro at the H19/Igf2 site. The model is quite robust to parameter choices with a broad peak of high performance in the range of *w* (2~4) and 〈*K_d,i_*〉/[CTCF] (5~10) (Fig. 2d). Also, the optimal parameters derived from training on GM12878 and HeLa are very similar (Supplementary Fig. 3a-c). The optimization curves for 5-fold cross validation are shown in Supplementary Fig 3d-f.

**Figure 2.**
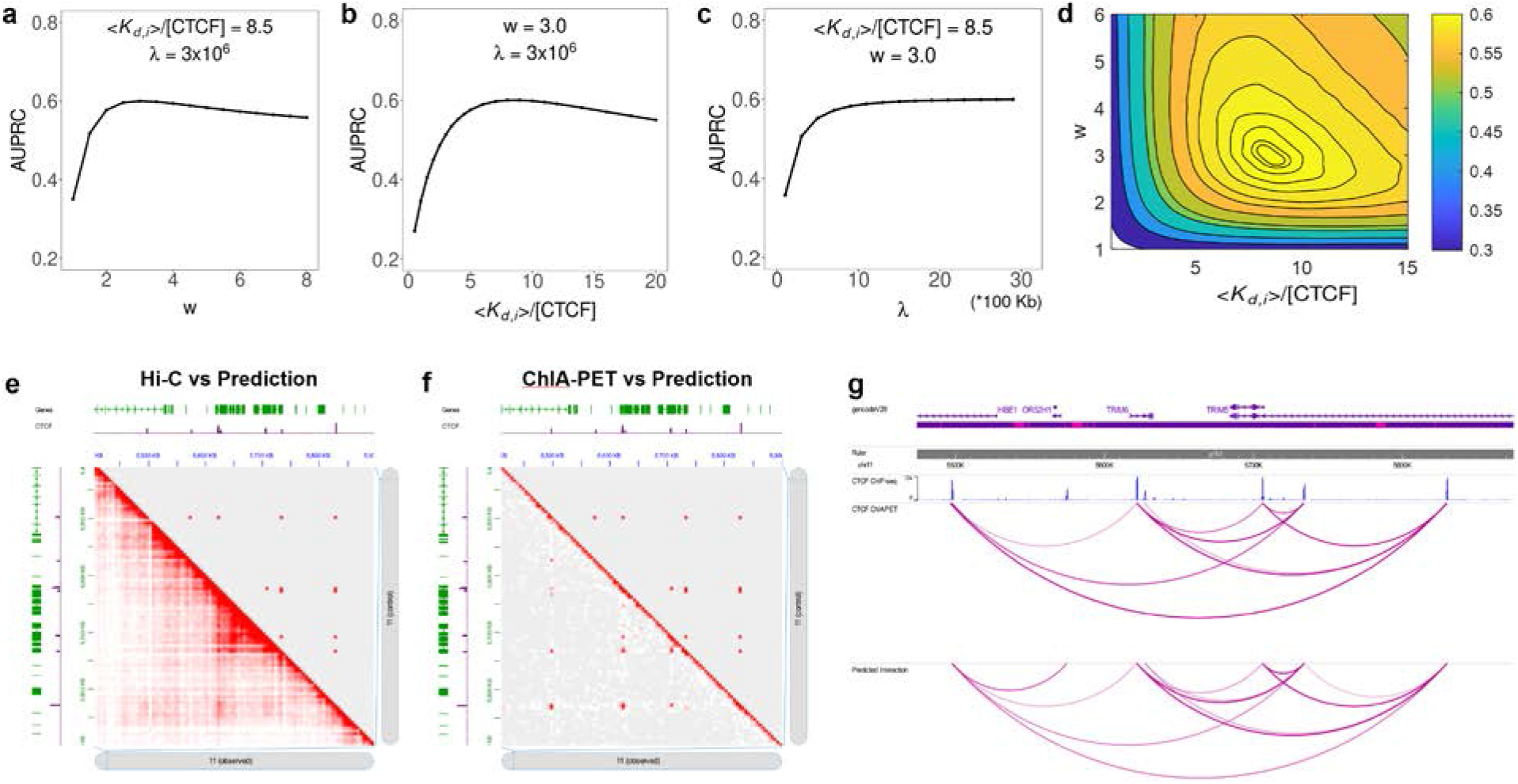
Model predictions compare favorably with Hi-C and ChIA-PET data in the TRIM5/6 locus. (a)-(d) Model performance is evaluated by area under precision-recall curve (AUPRC) as parameters are varied individually (a)-(c), and by grid search (d). Model predictions (upper-right) compared to (e) Hi-C data (bottom left) and (f) ChIA-PET data (bottom left). CTCF ChIP-seq signal is also shown as purple tracks in (a,b). The model predicts most of the direct CTCF ChIA-PET loop interactions, and Hi-C picks up additional contacts within the loops (or TADs) that are not the result of direct CTCF-CTCF interactions. (g) In the same locus, loops called by our model and ChIA-PET data are quite similar, as visualized using the WashU epigenome browser.

For λ, we expected the optimal value to be around the average loop length of 300kb, as reported in previous literature^1,2^. However, the agreement between our model and the ChIA-PET data increases monotonically with λ, which implies that distance information is dispensable for the prediction of CTCF interactions, as larger λ reduces the variation of the exponential term with distance (Fig. 2c). Moreover, leaving the distance-associated exponential term out completely makes the agreement with data slightly better. This stands in contrast with the general view that distance regulates chromatin interaction frequency. Previous Hi-C studies have reported a power law decay relationship between chromatin contact frequency and genomic distance^12^. For completeness, we also compared a power law decay with exponential decay, replacing *D_ij_* with 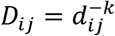, but performance was slightly degraded for all choices of *k* relative to the exponential distance function (Supplementary Fig 4(c)-(f)). The distribution of CTCF loop lengths is constrained by the genomic position of CTCF binding sites, and unlike Hi-C interactions does not follow a power law distribution, but the loop length distribution of our predictions is in close agreement with the measured length distribution from Fig S2.g from Ref. ^16^ (as shown in Fig. 2h). We shall directly address the apparent paradox that predictive performance is independent of loop length in detail below.

### Loop competition and extrusion model accurately predicts formation of CTCF-mediated loops

We applied our quantitative model of loop competition and extrusion (Eq. 5) to CTCF ChIA-PET data to predict CTCF interaction specificity. A total of 55,189 and 21,560 significant interactions with CTCF binding both anchors are identified for GM12878 and HeLa. All ChIA-PET detected CTCF-mediated loop interactions were labeled as positive samples, and all other (non-interacting) CTCF pairs within 1Mb were labelled as negative samples. Due to different sequencing depth and cell-type variability, the positive versus negative class ratio is roughly 1:20 for GM12878 and 1:37 for HeLa, with non-interacting CTCF pairs far outnumbering interacting pairs. A small fraction of loops had more than one CTCF binding peak at one of the anchors, when these could not be unambiguously assigned they were removed from the analysis.

In addition to the systematic performance evaluation by AUPRC described below, one specific example comparing our model predictions with Hi-C^6^ and ChIA-PET data is shown in the TRIM5/6 locus in Fig. 2e-2g. In this locus our model predicts a complex pattern of CTCF interactions that closely matches the ChIA-PET interaction counts. Hi-C picks up additional interactions within each CTCF loop which are not directly due to CTCF-interactions. Two other genomic loci are compared in Supplementary Fig. 5.

To assess the importance of each feature in our model, we trained on each individual feature and all combinations of features, including: CTCF binding intensity, CTCF motif orientation, distance and loop competition. An interaction probability *p_ij_* was predicted for all positive and negative pairs for each model, and was then compared to the true class label. Due to the huge class imbalance of CTCF interaction datasets, we employed area under the precision-recall curve (AUPRC) to evaluate model performance (see Method). For both GM12878 and HeLa cell lines, we observed that none of the four features alone could accurately predict interaction specificity of CTCF (AUPRC 0.2~0.3) while combining them increased the performance significantly (Fig. 3a-3b, Supplementary Table. 1). The best performance is given by the complete model, combining CTCF binding intensity (BI), CTCF motif orientation (Ori) and loop competition (LC), with AUPRC = 0.601. Performance on cross-fold validation test sets was: AUPRC=0.6005, std = 0.003.

**Figure 3.**
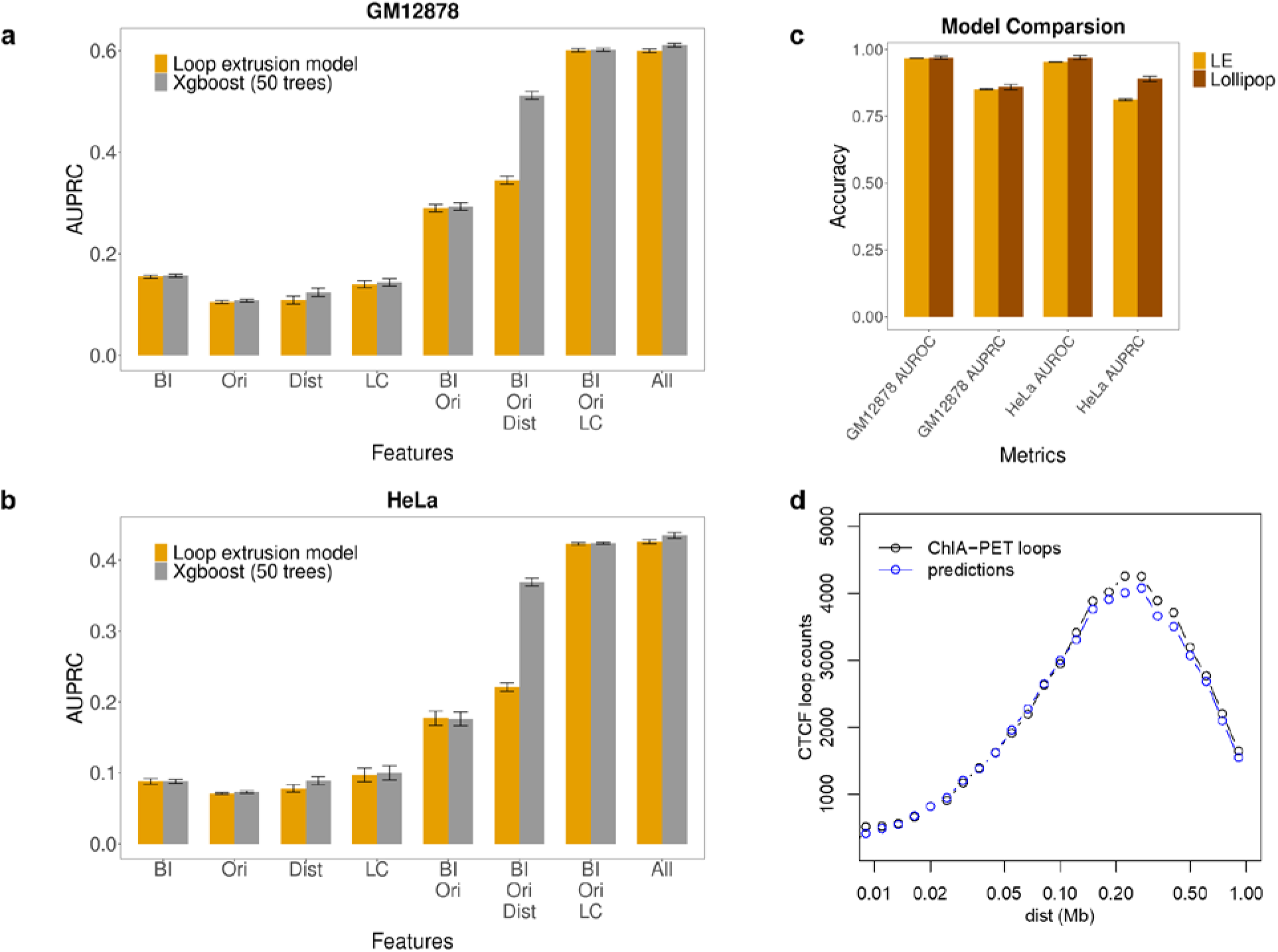
Model performance evaluation and feature importance. (a)-(b) Performance of our model with different combination of features for GM12878 and HeLa. BI – CTCF binding intensity; Ori – CTCF motif orientation; Dist – distance; LC – loop competition. Performance is also compared against xgboost model with 50 trees. Down sampling of 10% of the data was repeated 10 times and 95% confidence intervals are shown. (c) Model comparison against Lollipop under class ratio 1:5 (positive vs. negative). (d) Loop length distribution for measured ChIA-PET loops and predicted interacting loops are quite similar.

This model combines these features in a functional form specific to an underlying mechanism of loop formation. To test this mechanistic assumption, we also constructed a more general machine learning model using boosted trees, with exactly the same features, to compare with our model. Surprisingly, the boosting model, with no constraints on the form of the nonlinearity among features, is only marginally better than our model (AUPRC = 0.602). This comparable performance increases confidence in the validity of the mathematical formulation of our model and the loop extrusion hypothesis. Furthermore, adding distance (Dist) as a feature does not significantly increase performance in either our loop extrusion model or the boosting model (AUPRC = 0.611). This confirms our earlier observation (from the insensitivity of performance to λ) that distance is weakly informative and seems to be redundant for our model in this task. Notably, even without distance information, the distance distribution of interacting CTCF pairs predicted from loop extrusion model is still extremely close to experimental data (Fig. 3d) and matches FigS2g of Ref ^16^. Results in HeLa cell line (Fig. 3b) are qualitatively consistent with GM12878, with reduced AUPRC attributable to the larger HeLa class ratio difference. We then compared our model with a previously published machine learning model, Lollipop, which successfully predicted CTCF-mediated loop with 77 different sequence and epigenomic features (Fig. 3c). Under the same class ratio 1:5, we found that in both cell lines, our loop extrusion model is nearly as accurate as Lollipop in terms of both AUROC (area under the receiver operator characteristic curve) and AUPRC, which indicates that the information contained in our model is quite comprehensive, relatively more compact, and more easily interpretable.

To evaluate the quantitative predictions of our model, we compared the predicted interaction probability of CTCF pairs, conditioned on their quantitative labels, to the PET counts from the ChIA-PET experiment. The model probabilities are highly correlated with PET count (C = 0.686 for GM12878 and 0.531 for HeLa) (Fig. 4a). In addition, positive and negative CTCF pairs are clearly separated by predicted interaction probability (Fig. 4b).

**Figure 4.**
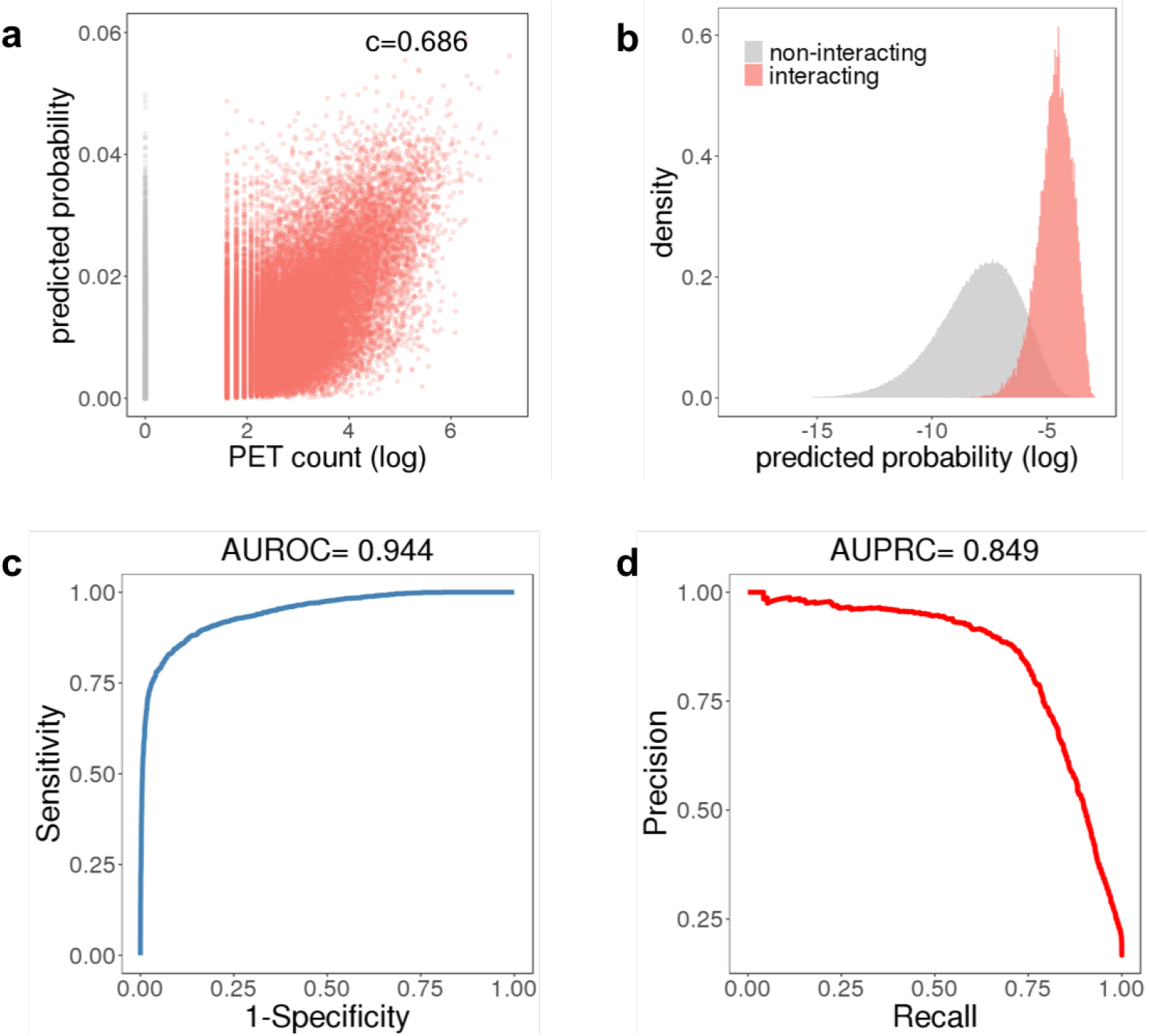
Model validation by quantitively assessing CTCF ChIA-PET and Micro-C dataset. (a) Distribution of PET count (log scale) against loop extrusion model predicted interaction probability. Red dots are interacting CTCF pairs while grey dots are non-interacting CTCF pairs. (b) Distribution of loop extrusion model predicted interaction probability. (c)-(d) Validation of model prediction performance on Micro-C CTCF loops with AUROC and AUPRC.

To validate our model on an additional external dataset, we predicted CTCF loops identified from a recently published high resolution Micro-C dataset.^36^ In total, 15,945 significant loops at 1kb resolution were detected in this dataset with HICCUPS.^6^ For purposes of predicting CTCF-mediated loops, we sampled positive loops with CTCF binding at both ends, and generated a five times larger negative set by sampling from non-interacting CTCF pairs. We applied our model on this dataset and achieved (AUROC = 0.944, AUPRC = 0.849) (Fig. 4 (c)-(d)), indicating that we are able to accurately predict CTCF interaction at a similar performance to those detected by ChIA-PET. Taken together, the analysis of CTCF ChIA-PET and Micro-C data shows that CTCF interaction can be successfully predicted from the loop extrusion model, and only requires information of local CTCF binding intensity, CTCF motif orientation and loop competition throughout the local neighboring region (up to 3Mb). We tested adding additional features to the boosting model, e.g. Cohesin ChIP-seq and DNase-seq signal, but found that these did not improve performance significantly (Supplementary Fig.1, Supplementary Table. 2).

### Loop competition is a more powerful predictor than distance

Because of the simple formulation of our model, we can evaluate the relative importance of each component to the loop formation process. First, we calculated the correlation between all pairs of features and PET count (Fig. 5a-5b). The only two features highly correlated with each other are distance and loop competition (Dist and LC). This correlation is to be expected, because the more distant two CTCF binding sites are, the more likely the existence of a competing loop becomes. But which of these correlated features is more predictive of CTCF interactions by itself, distance or loop competition? Almost all studies of genome-wide chromosomal conformation capture experiments, including Hi-C, ChIA-PET and Micro-C, have reported that a longer distance between two regions is associated with reduced interaction frequency^6,16^. Intuitively, distant regions contact less frequently by diffusion in three-dimensional space, but the precise mechanism of the observed loop distance dependence has not yet been supported by much direct experimental evidence. It is possible that the distance dependence is associated with some other factor which determines loop formation.

**Figure 5.**
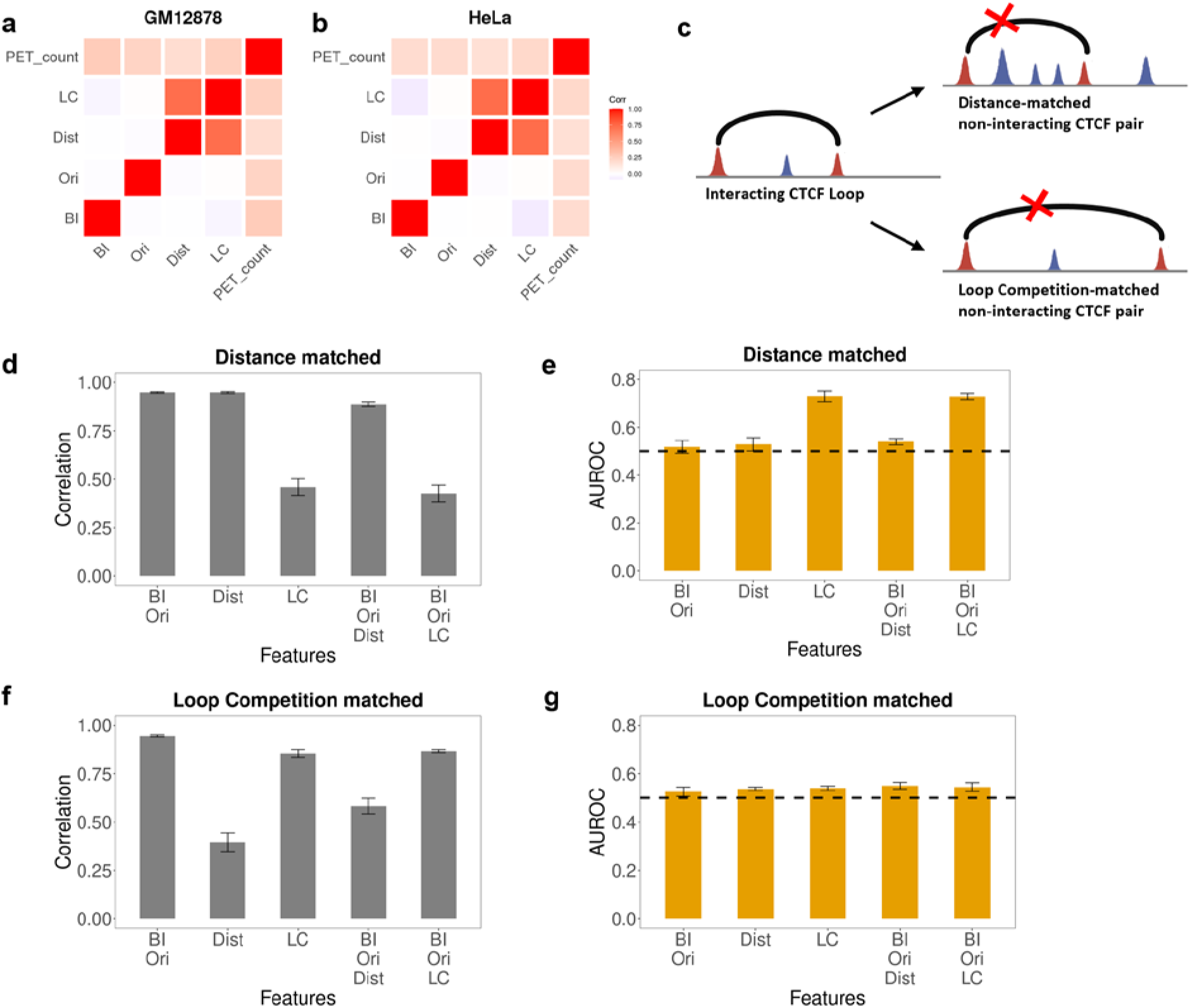
Loop competition is a more crucial determinant than distance. (a)-(b). Correlation of features (CTCF binding intensity, CTCF motif orientation, distance, loop competition (LC) and PET count) across all positive and negative pairs (log scale). Since loop competition and distance are correlated, we designed an additional experiment to isolate their relative informative value. (c) We generated distance-matched and loop competition-matched subsets of the full data by choosing a negative pair (marked with X) for each positive pair with either LC or distance matched within a factor of two. (d),(f). Correlation between positive and negative set for different combinations of features in both matched settings. (e),(g). AUROC of loop extrusion model with different combinations of features in both settings. Down sampling of 10% of the data was repeated 10 times and 95% confidence intervals are shown. Since LC adds informative value in a distance matched evaluation set but the converse is not true, loop competition is the more predictive feature.

To determine the relative importance of distance and loop competition, we generated distance-matched and loop-competition-matched test sets by sampling the ChIA-PET data to isolate the contributions of each feature (Fig. 5c). In distance-matched sampling, for each positive loop, we selected one negative loop with similar CTCF binding intensity, CTCF motif orientation, and distance (within a factor of two for BI and Dist) (see Methods). In other words, every feature except loop competition is matched between this negative set and the positive set. Compared to the full dataset, it should be harder to distinguish the positives and negatives in this set because loop competition is the only unmatched feature. By evaluating our model with on this distance matched set with different subsets of features, we find, as expected, CTCF binding intensity, CTCF motif orientation or distance are not useful for prediction on this subset (Fig. 5e). In contrast, the model including loop competition (LC) reached AUROC = 0.730, indicating that loop competition alone is predictive in this context and carries unique information about loop formation that doesn’t exist in distance alone. We next generated a loop-competition-matched sample in a similar fashion, selecting positive and negative loops with similar levels of loop competition (within a factor of two) but unmatched distance, in which loop competition for each CTCF pair is determined by Eq. 4. In contrast to the distance matched subset, in the loop-competition-matched subset, distance is not predictive of CTCF loop formation, showing that distance itself cannot explain CTCF interaction specificity (Fig. 5g). The fact that loop competition is predictive in a distance matched context, while distance is not predictive in a loop-competition-matched context, indicates that loop-competition is the more informative feature. This test suggests that distance can be a predictive feature because it can serve as a proxy for loop competition when loop competition is not an explicit feature of the model. Our results show that the negative correlation between distance and contact frequency is likely to be mediated by the effect of loop competition. Consistent with this interpretation, distance has the weakest correlation with the PET count of loops among the four features (Supplementary Fig. 6). These computational experiments confer support for loop competition as an important determinant of CTCF interaction specificity.

### Testing loop competition by CTCF disruption in population Hi-C data

Our model makes quantitative predictions about how a single CTCF binding site disruption would be expected to impact the interaction strength of multiple CTCF loops in a genomic locus. Since loop competition is a dominant feature in our model, attenuation of one loop would in turn facilitate or strengthen flanking and overlapping loops. Specifically, our model predicts that if a given CTCF binding site is disrupted by sequence variation or mutation, it will be less likely to form a loop^17^, and consequently other CTCF pairs spanning the disrupted site would be more likely to interact, as a result of reduced loop competition. A previously published dataset which measured Hi-C loop interaction frequencies in lymphoblast cells derived from 20 individuals provides a direct means to test our model predictions of how CTCF disruption affects loop strength^37^. Natural genetic variation in this sample disrupted 49 CTCF binding sites by SNPs. For each CTCF binding site disruption, we separated individuals into two groups (strong or weak CTCF motif, as strong motif defined as those consistent with CTCF PWM at key positions in dashed boxes in Fig. 6a), and calculated the ratio of average contact frequency in 40kb bins in neighboring 800kb windows in the two groups (Fig. 6a). After aggregating this data for all 49 CTCF sites, we observed that on average, bins that represent interactions between pairs of loci that span the CTCF motif (labeled as ‘Cross’) exhibit a higher normalized interaction frequency in weak vs. strong motif individuals (100/100 bins higher for weak motif individual), consistent with reduced loop competition in our model (model predictions shown in Fig. 6b). In addition, interactions that do not span the CTCF binding site (labeled as ‘Outside’) have much weaker differences, and their direction of change is much more random (52/90 bins higher for weak motif individual). This data supports the role of loop competition in loop formation and provides an interesting mechanism of how genetic variation could affect chromatin conformation. It is also consistent with a recent report that subtle quantitative changes in CTCF loop strength could lead to phenotypic variation in gene expression^38^.

**Figure 6.**
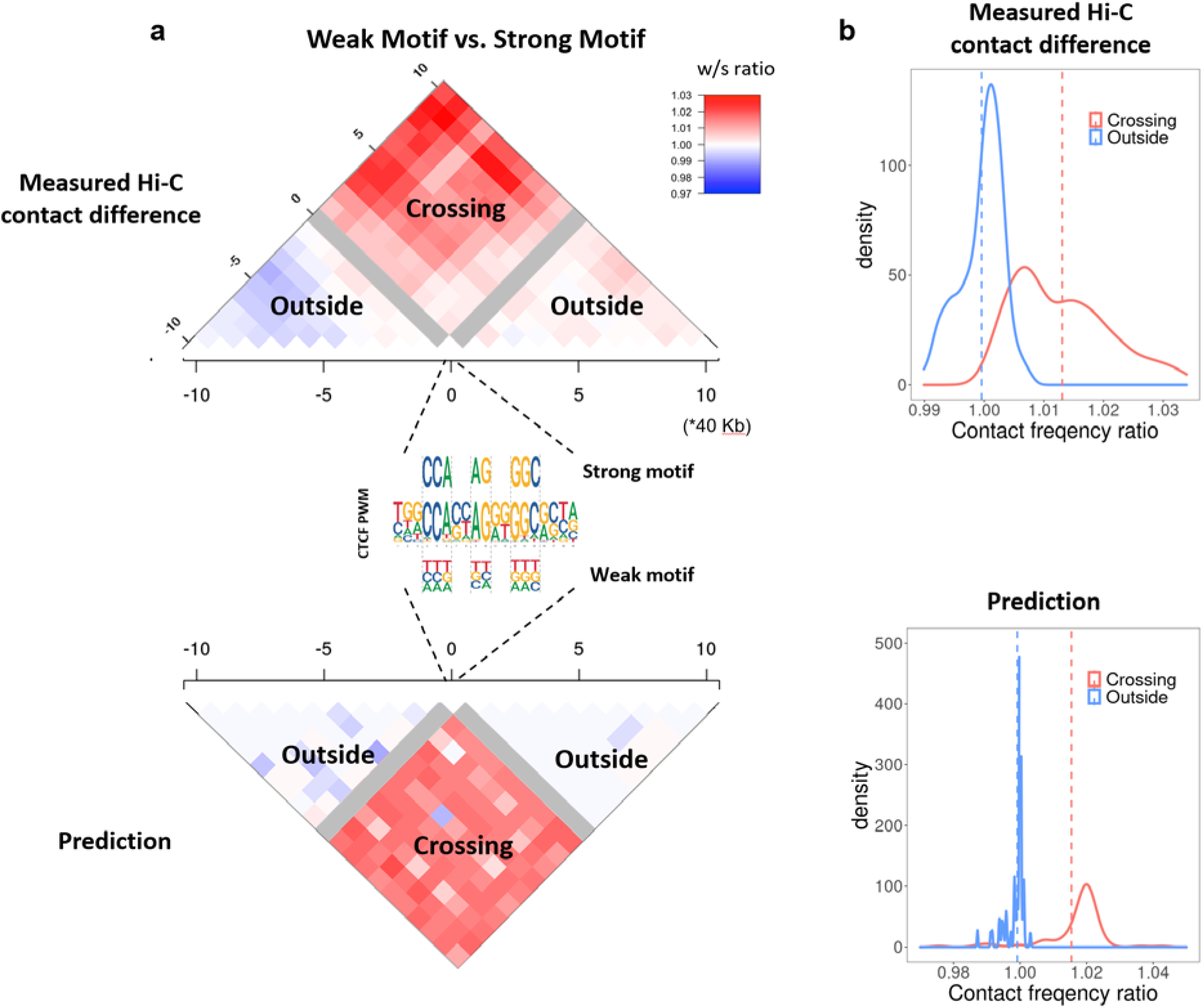
Loop competition predictions are consistent with changes in chromatin interaction frequency induced by naturally occurring CTCF binding site disruption. (a) Measured differential Hi-C contact frequency flanking SNP disrupted CTCF sites. The contact ratio for weak vs. strong CTCF motif genotype in a population of 20 individuals^37^ is shown. The heatmap is partitioned into 40kb bin pairs. Gray bins directly overlap the disrupted CTCF binding site on one end, and loops which span the CTCF motif (Crossing) or do not span the CTCF motif (Outside) are indicated. Only loops which span the disrupted CTCF motif have increased contact frequency (top, red), consistent with reduced loop competition from our model predictions (bottom). Only SNPs which disrupt the indicated informative positions in the CTCF motif are used, and the strong or weak versions are labeled on the PWM from Jaspar MA0139.1. (b) The same data is used to generate the contact frequency ratio distribution for the two classes (Crossing and Outside) of bin pairs for measurements (top) and our model predictions (bottom).

### Loop extrusion model predicts effect of CTCF binding perturbation and WAPL knockout

Many in vivo perturbation experiments have been carried out to study the role of CTCF in loop formation and gene regulation^39^. In addition to knocking out CTCF, many studies have deleted or inverted the CTCF binding motif, revealing a great preference of convergent CTCF motif orientation for chromatin loops^17,19,40^. These studies provide important additional contexts to test our model. In one particular study, the effect of CRISPR targeted deletion or inversion of a CTCF binding motif in mouse embryonic stem cells (mESC) was measured with 4C^17^. To make predictions in the three loci tested, we used CTCF ChIP data measured before and after the perturbation, modified *w* for inversions, and we calculated the corresponding loop interaction probability from our model. Before CRISPR editing, the predicted interaction probabilities matched the 4C loop measurements very well (Fig. 7a-7c, only the strongest 4C loop corresponding to the target site is shown). Moreover, after CRISPR editing, our model successfully predicts the loss of the wild-type loop induced by both deletion and inversion of CTCF binding motif for Malt1, Sox2 and Fbn2 loci (Fig. 7d-7f). Although inversion of the CTCF binding site does not change CTCF binding dramatically, inversion affects loop formation through the parameter *w*, and the reduced interaction probability is consistent with the observed reduction in 4C signal.

**Figure 7.**
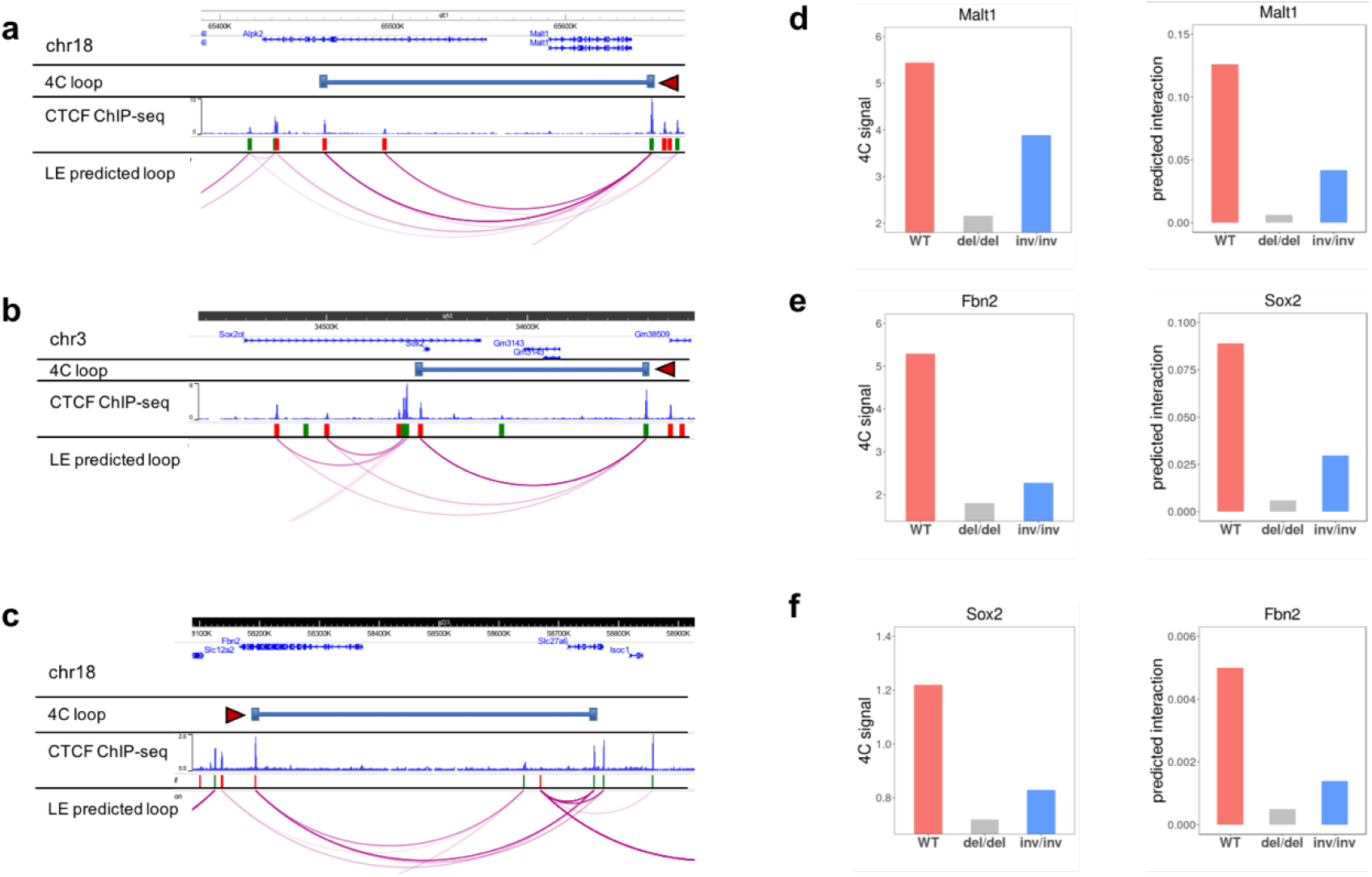
Loop extrusion model predicts the effect of targeted CTCF disruption and inversion on chromatin interactions. (a)-(c). Comparison of contact profiles of 4C-seq measurements and our loop extrusion model at the Malt1, Sox2 and Fbn2 loci. Only the strongest loop of the targeted CTCF binding site (indicated by dark red triangle) from 4C-seq is shown. The orientations of flanking CTCF motifs are indicated by red (forward) and green (reverse) bars. Our loop extrusion model predicted interacting CTCF pairs are shown, with darker color corresponding to higher interaction probability. (d)-(f). 4C-measured interaction frequency and loop extrusion model predicted probability of looping for wild-type and after CRISPR deletion or inversion of the targeted CTCF binding site.

Alternatively, the activity of Cohesin can be modulated through the Cohesin unloading factor WAPL^41,42^. It has been reported that upon WAPL knockout the overall chromatin structure transforms into a more condensed state, with an increase in loop number and size. Although it is known that WAPL knockout increases Cohesin residence time on chromatin^34^, the means by which this changes loop interactions under the same set of CTCF boundary locations remains unclear. Since our original model was derived under the normal assumption of constant WAPL activity, we modified our model slightly to predict the effect of WAPL knockout on CTCF-mediated loops. In this WAPL-KO modified model (see Supplementary Fig. 7a and Methods), following previous work^11,12,29^, we assume that Cohesin is not completely blocked at CTCF loop anchors, but can pass through with some small probability, *s*. WAPL knockout increases the residence time of Cohesin, which consequently has a greater chance of passing through boundary CTCFs. With enhanced pass-through probability, the effect of loop competition is reduced because Cohesin is moving more freely in this case. Through testing the WAPL-KO corrected model, we found pass-through probability is positively correlated with total loop number and average loop size. At pass-through probability around 0.4, we faithfully reproduced experimental results from WAPL knockout in HAP1 and Hela cell lines^41,42^ (Supplementary Fig. 7b-c). We also compare Hi-C data to model predictions in the context of the WAPL knockout in Hela in Supplementary Fig. 7d.

### CTCF loops constrain enhancer-promoter interactions

An important proposed function of CTCF loops is to shape local chromatin architecture to constrain interactions between other types of regulatory elements, especially enhancers and promoters^43^. According to this idea, enhancer-promoter interactions should preferentially occur within CTCF loops, and not to cross CTCF loops. To assess this hypothesis with our model, we took an integrated enhancer perturbation dataset consisting of 4194 enhancers and 65 gene promoters in the K562 cell line from 11 studies^44–54^. We counted the number of CTCF loops crossed by each enhancer-promoter (E-P) link and the number of CTCF loops which contain each E-P link. We then compared the fraction of interacting vs. non-interacting E-P pairs in loop-crossing and loop-containing events. Consistent with our hypothesis, based on K562 CTCF ChIA-PET measured loops, we observed a 2.9-fold enrichment of true E-P links in the group that does not cross any CTCF loop, compared to the group crosses one or more CTCF loop. (Fig. 8a-8b) Similarly, there is a 1.6-fold enrichment of true E-P links in the group that is contained by one or more CTCF loop, compared to the group that is not contained within any CTCF loop. Strikingly, the level of enrichment of ‘not cross’ and ‘contain’ groups increased dramatically to 6.6 and 7.8, using our loop extrusion model CTCF loops instead of ChIA-PET annotated loops. Although this clearly lends support to our model, it may seem perplexing that a model trained on ChIA-PET data seems to be more consistent with expectations of E-P loop crossing than the ChIA-PET data itself. One possible explanation is that our model prediction is largely coming from CTCF ChIP-seq intensity, orientation, and loop competition, all single-point measurements, while ChIA-PET interactions are pairwise and require much more sequencing depth to achieve comparable signal-to-noise ratios. Technical considerations may contribute to false positive or negative loop interactions in the ChIA-PET data which do not constrain E-P interactions as effectively as those predicted by our model. While genomic ChIA-PET data with thousands of loops can reliably determine the parameters in our model, the model may actually be more accurate at predicting functional CTCF loops in a given locus.

**Figure 8.**
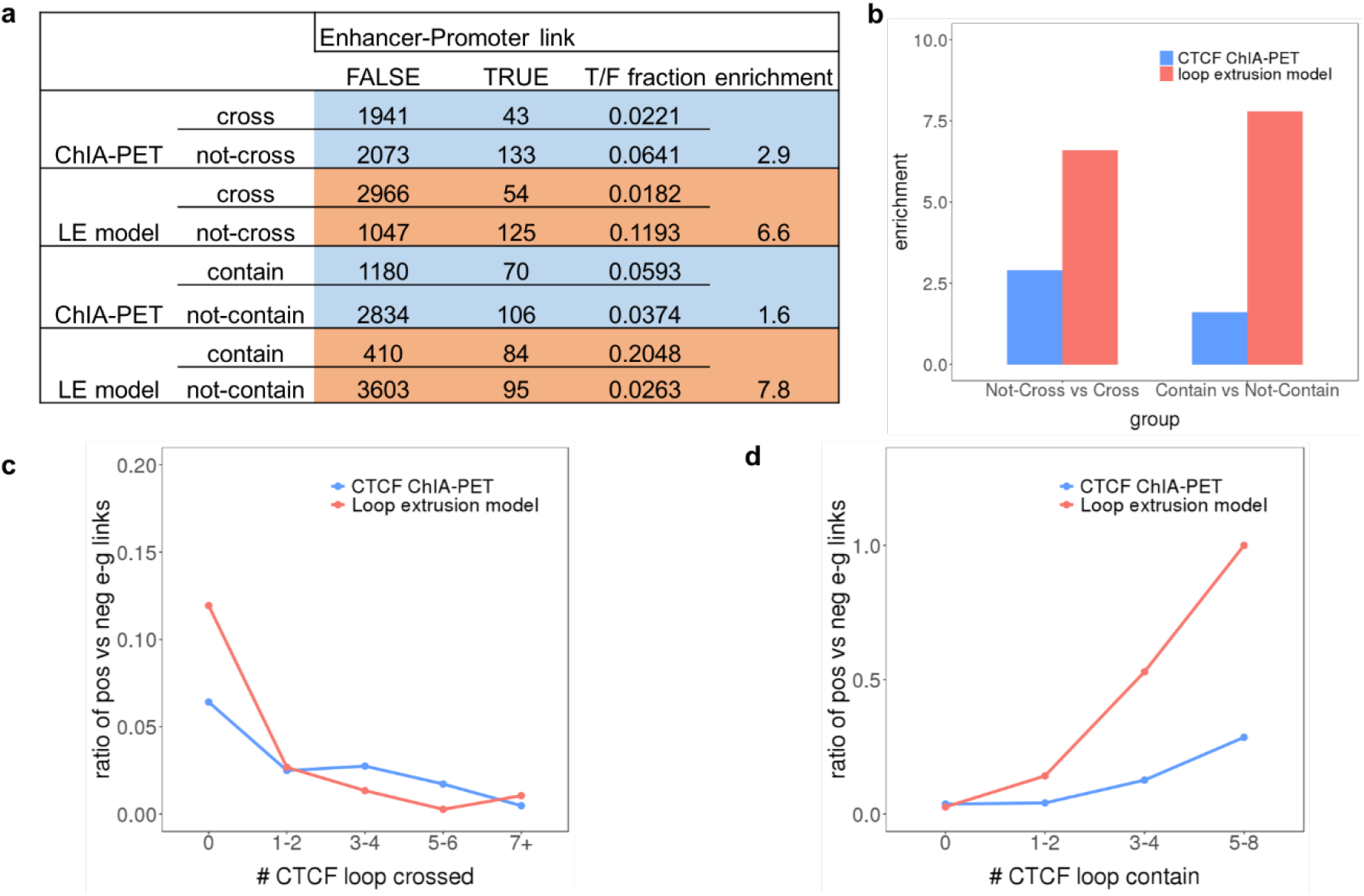
CTCF loops are predicted to constrain enhancer-promoter interactions, but loop extrusion model predicted loops do so more accurately. (a) Counts of true (interacting) and false (non-interacting) enhancer-promoter (E-P) pairs according to whether they cross, or are contained within CTCF loops. Ratios of true and false E-P links are also shown (T/F). (b) Enrichment of T/F ratio between each group are calculated and compared between CTCF ChIA-PET annotated loops and loops predicted by our loop extrusion model. Strikingly, the predicted CTCF loops are much more enriched for loops which contain (and do not cross) E-P interacting pairs. (c) T/F ratio against the number of CTCF loops each E-P link crosses is plotted. (d) T/F ratio against the number of CTCF loops containing each E-P link is plotted.

### CTCF binding intensity is predictive of cell-type specific loops

Next, we investigated the cell-type specificity of CTCF loops and whether cell-type dependent CTCF loops could be predicted by the loop extrusion model. Cell-type specific chromatin interactions are of great interest because they have been demonstrated to be an important mechanism for gene regulation in lineage differentiation^43,55^. We noticed that GM12878 and HeLa ChIA-PET experiments have very different numbers of detected loops, but this is mostly due to differences in sequencing depth. To eliminate this bias, we constrained our analysis to the strongest 10,000 CTCF loops in each cell line. We find that these top loops are quite conserved. Over 75% of them are shared between the two cell lines (Fig. 9a-9b). These cell-type specific CTCF loops can also be predicted with our loop extrusion model, because the difference in their activity is strongly associated with CTCF binding intensity in GM12878 vs. HeLa (AUPRC 0.955 for GM12878-specific loops, 0.739 for HeLa-specific loops) (Fig. 9c-9d).

**Figure 9.**
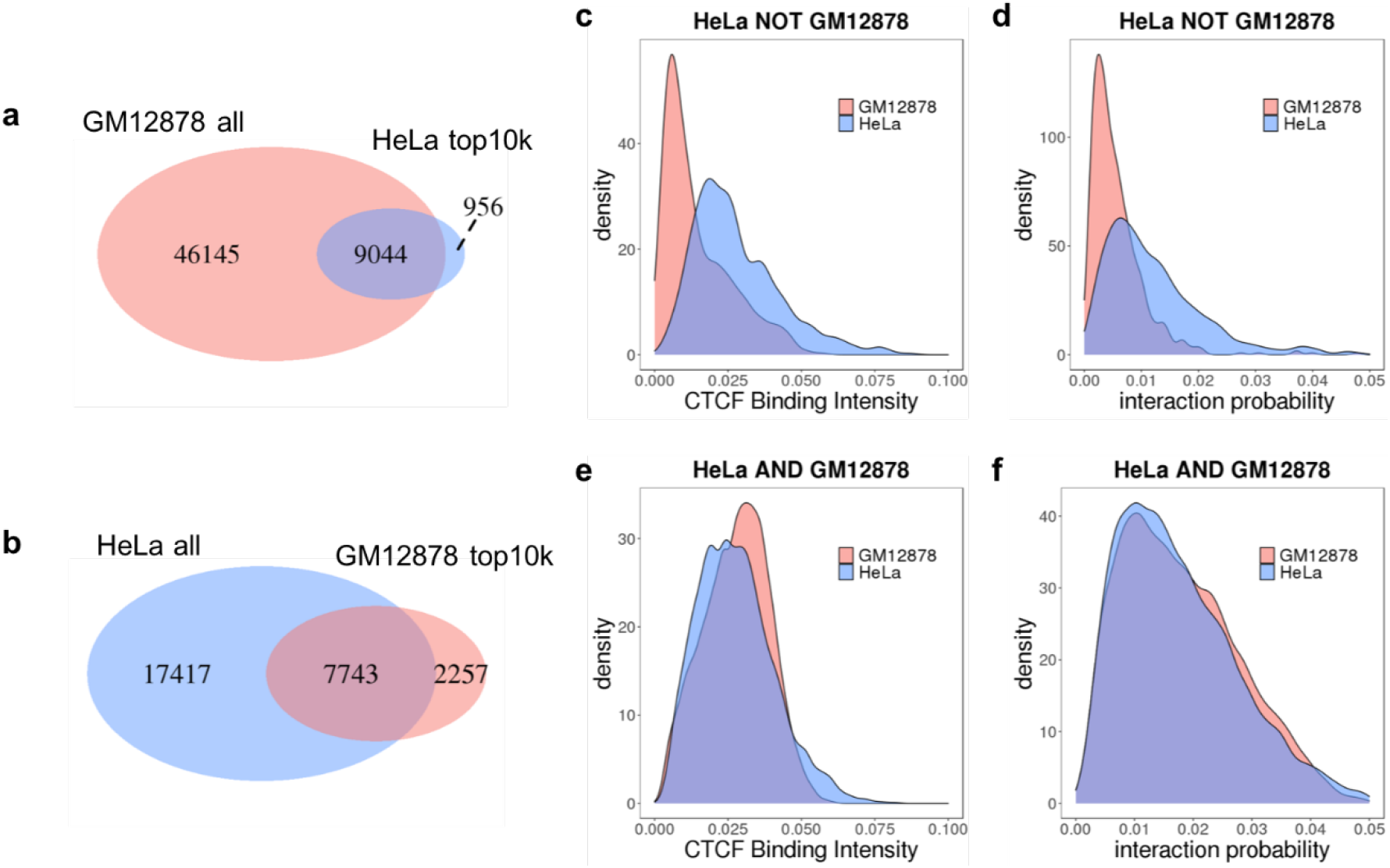
CTCF binding intensity is predictive of cell type specific loops. (a)-(b). Venn diagram of CTCF-mediated loops identified from GM12878 and HeLa ChIA-PET. Only the strongest 10,000 loops are compared against each other due to different sequencing depth. (c)-(f) CTCF binding intensity distribution and predicted interaction probability distribution for HeLa specific CTCF loops and shared loops.

## Discussion

Recent progress in 3C techniques has enabled comprehensive annotation of higher order chromatin architecture, including CTCF-mediated loops. Predicting CTCF-mediated loops is a crucial first step toward understanding the mechanisms controlling regulatory element interactions and transcriptional regulation. While dramatic progress has been made mapping regulatory element activity in a large collection of cells and tissues^56^ and detecting active TF binding sites in these elements with machine learning^57^, connecting regulatory element activity to dynamical models of gene networks and cell state transitions is in its infancy^57,58^, in large part due to our limited understanding of what controls enhancer-promoter interactions and how competitive or cooperative interactions between multiple enhancers are integrated at a target promoter. It has been shown that the interaction between enhancers and their target gene promoters cannot be predicted solely from local epigenetic signals^59,60^. The missing element is very likely to be the spatial organization of chromatin, as disruption of CTCF-mediated loops have been confirmed to be able to change the expression of genes both inside and outside of the loop. Moreover, recent sequence-based modeling of enhancer-promoter interactions have also identified CTCF binding as the most important player^61^. We were motivated to develop a simpler model of CTCF interactions after a machine learning approach showed that CTCF interactions in ChIA-PET data could be predicted with high accuracy using a large set of epigenomic features^26^.

Our model correctly distinguishes interacting CTCF pairs from a vast number of non-interacting CTCF pairs. This could not be achieved using only convergent CTCF motif orientation as a feature, as many convergent CTCF motifs do not interact, and some true interactions are tandem. Our model is easily interpretable, as the contribution of each component is independently modelled by its corresponding probability. We validate our model on a wide range of complementary datasets: ChIA-PET, Micro-C, Hi-C, genetic variation in CTCF binding sites, CRISPRi perturbation of loop anchor binding sites, and by the predicted CTCF loops’ ability to constrain enhancer promoter interactions.

Our analysis reveals that the distance between two CTCF pairs, previously thought to be important for constraining chromatin interactions, actually becomes unimportant when we explicitly calculate the contribution from loop competition. This raises the question of whether this is specific to CTCF-mediated loops or a broader class of 3D chromatin interactions. A recent study from E.coli proposed an interesting ‘small world’ hypothesis that because the bacteria genome is so small and compact, different parts of the genome, regardless of their linear position, are all equally likely to randomly collide with each other^62^. This is unlikely for the human genome given its huge size and partitioning into chromosomes, but may be true within single TADs.

The concept of loop competition arises naturally from the loop extrusion process (Fig. 1b). The loop competition hypothesis is that CTCF pairs across an existing loop are less likely to be formed, while those within or outside it are unaffected. This idea is supported by observations that strong CTCF corner peaks prohibit cross TAD interactions^5^. Disruption of CTCF binding sites and rearrangement of corresponding CTCF loops facilitates ectopic interactions between enhancers and gene promoters over long distances and could potentially give rise to severe pathogenic phenotypes like polydactyly^8^. We used our quantitative predictions of loop competition to predict the consequences of CTCF motif sequence variation on neighboring chromatin interactions, and showed that the impact is significant, consistent with our modeling, and detectable over several hundred kilobases. Importantly, this result shows that chromatin architecture should not be viewed simply as a combination of independent structural units, since there can be extensive interplay between adjacent elements.

We found that CTCF-mediated loops are rather stable across cell lines, consistent with previous studies^55^. However, although less common, when cell-specific CTCF loops do occur, they can be consequential, as cell-type specific loops are often accompanied by gene activation or repression^55^. Our modeling shows that these cell-specific CTCF loops are mediated by variable cell-specific activity of CTCF binding sites.

Although our model is trained on ChIA-PET data collected from a population of cells, the probabilistic formulation of our model is consistent with quantitative measurements of the number of CTCF molecules in a single cell^33,35^, which suggest that not all CTCF binding sites detected by ChIP-seq are consistently occupied, but that CTCF and Cohesin are popping on and off the genome as extrusion occurs. This is supported by the probabilistic form of our loop competition term. While in any given cell, an extrusion through a given CTCF binding site may or may not be blocked, the time or population average of the probability of loop formation is what correlates with ChIA-PET contact frequency.

It is worth noting that the model presented here does not rule out other chromatin organization or loop formation mechanisms. For example, emerging experimental and computational evidence has suggested that phase separation could be the underlying mechanism for the larger scale A/B compartmentalization observed in Hi-C contact maps^21,63,64^, and we envision the Cohesin/CTCF loop formation described here as operating on a shorter length scale within compartments. In addition, although Cohesin degron experiments provide compelling evidence for a model where Cohesin extrusion and CTCF blocking is the primary determinant of CTCF-mediated loop formation, this is not the only possibility. Loops could also be formed by interactions between other protein bound complexes (e.g. enhancers and promoters) modelled by “Strings-and-Binders” or SBS polymer models^24^ that may dominate within CTCF loops. These polymer physics based models have been used to predict impact of structural variants on 3D structure^22,65^, and to predict contact frequency variability between individual cells^21,66^. Our model can make a subset of these predictions, but our current formulation focuses only on CTCF mediated loop interactions.

In summary, we constructed a mathematical framework to predict single loop level chromatin architecture based on a loop extrusion model. We validated our model by showing that the model predictions are in agreement with four diverse experimental datasets, which in turn provides substantial support for the loop extrusion hypothesis. Although we have extensively tested our model on existing data, prediction of CTCF looping interactions in blind computational assessment challenges such as CAGI^67^ would be an interesting next step, as these efforts are beginning to focus more on regulatory processes^68^. We expect our loop extrusion model to be useful for further exploration of both the features and mechanisms of chromatin packaging and its impact on gene regulation, and as a component of more comprehensive models of enhancer-promoter interactions.

## Methods

### Loop competition and extrusion model

The loop extrusion model is a hypothesis that describes the formation of CTCF-mediated loops via Cohesin movement. The probability of CTCF loop formation is determined by four components.

#### 1) CTCF binding intensity

The occupancy of CTCF is characterized by the standard calculation of chemical equilibrium,^28^

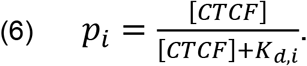

[CTCF] corresponds to the concentration of CTCF, and is represented by the normalized read count in the window of the binding site. *K_d,i_* is the equilibrium dissociation constant for each CTCF binding site. This dissociation is not necessarily simply due to the strength of the CTCF binding motif, as local chromatin context and interactions with flanking factors may contribute to CTCF binding. Therefore we will estimate this local *K_d,i_* from the CTCF ChIP-seq signal. We can combine the unknown *K_d,i_* and [CTCF] to write 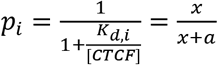, and we will further assume that the local ChIP-seq signal *x* is inversely proportional to *K_d,i_*/[CTCF], with a scaling factor of *a*. We will learn the best value of the parameter *a* from the ChIA-PET data. The precise form of the ChIP-seq signal scaling with 1/*K_d,i_* is not critical, as we have also tried a different parameterization of the binding probability using *p_i_* = tanh(*ax*), which yields almost equivalent performance(Supplementary Fig 4b).. With the assumption that CTCF binding at each site are independent, joint probability of CTCF binding at two sites at the base of a loop is given by their product *p_i_* ∙ *p_j_*.

#### 2) CTCF motif orientation

CTCF-mediated loops have strong motif orientation preference, with convergent motifs being the most favored configuration and divergent motifs being the least favored. To model this difference, we modeled the relative stability of convergent, tandem, and divergent loops as 1, 1/ *w*, and 1/ *w*^2^, where *w* is a scalar, *w* > 1. This can be interpreted as an orientation dependent stability of the CTCF-Cohesin complex at the base of a loop, where each “non-inward” CTCF motif decreases the stability the complex by a factor of *w*. We also tested a more general form of orientation dependent stability as 1, 1/ *w_1_*, and 1/ *w_2_* for convergent, tandem and divergent loops, and obtained very similar results (Supplementary Fig 4a).

#### 3) Distance

A strong anti-correlation has been found between chromatin contact frequency and the distance between the interacting regions in genome-wide 3C experiments. Various probabilistic distributions have been used to fit this relationship. In our model, this distance dependence could arise from a constant probability of cohesin dissociating from the chromatin fiber as it translocates over longer distances. A constant dissociation probability would lead to an exponential distribution (cumulative probability of staying on the fiber):

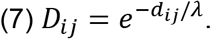

The average CTCF loop length would scale with this parameter λ. Previous Hi-C studies have reported a power law decay relationship between chromatin contact frequency and genomic distance at a population level^12^. This has also been observed in polymer physics based modeling^24^. The observed power law dependence of contact frequency arises from the genomic distribution of distances between loop anchors in the genome, and in our case depends strongly on the spacing between CTCF binding sites, and often averaging over many events leads to power law scaling. But for completeness, we also directly compared power law distance decay with exponential decay, by directly modeling *D_ij_* as:

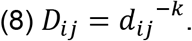

This led to slightly reduced predictive accuracy after fitting the parameter *k* on ChIA-PET data (Supplementary Fig 4c-f).

#### 4) Loop competition

The process of loop extrusion implies a competition between two Cohesins translocating along the same linear chromatin segment. Since the final state of the extrusion is Cohesin contacting a CTCF barrier pair, this further implies a competition between CTCF pairs which overlap each other. ‘Overlapping’ here is defined with regard to the window between CTCF binding sites. As Cohesin cannot move across another Cohesin on a pre-formed loop, a prerequisite of loop formation would be that no overlapping loops exist, therefore

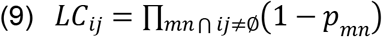

describes this probability. Because the loop competition term involves *p_mn_* but contributes to *p_ij_*, it should be calculated iteratively. We implemented an iterative solution with successive over-relaxation. If successive iterations are labelled by *k*, we used

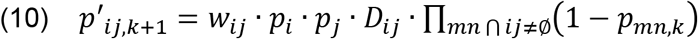

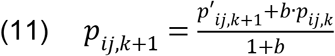

where *p_ij,k_* is the normalized looping probability between *i* and *j* at iteration *k*.

This scheme converged for a wide range of relation rates *b*. (Supplementary Fig. 4g). But we also noticed that the full iterative solution of Eq. 3 is consistent with a simpler method, which just requires that all CTCF sites internal to the loop *ij* are unoccupied, using *p_m_* from Eq. 1 for the probability of occupancy of site *m*. This closely approximates the probability that no overlapping loop exists and avoids the necessity for iterative solution. Thus in our evaluations we actually used an approximate loop competition term which requires that all CTCF sites between *i* and *j* (the current window) are unbound:

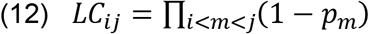

The performances of the iteratively trained model and approximation model are very close (AUPRC 0.589 vs 0.601 for GM12878, Supplementary Fig. 4g).

The final probability of loop formation is the joint probability or product of these five terms (two CTCF binding probabilities, one from each of the two sites), initially assuming they are independent.

### Model performance evaluation

We evaluated the model by defining a set of positive CTCF ChIA-PET loops (counts≥ 4 for GM12878, counts≥ 3 for HeLa) and a set of negative loops (pairs of CTCF binding events within 1MB not called as ChIA-PET loops). We removed ChIA-PET loops that did not map to a CTCF ChIP-seq peak, or did not have a strong enough CTCF binding site motif to unambiguously assign an orientation at either anchor. We then used the loop extrusion model *p_ij_* as a threshold variable for prediction to generate Precision-Recall curves and calculate AUPRC. We trained on both 5-fold chromosomal training-test splits and on the entire dataset with identical parameter optimization and performance (Supplementary Fig.3d-f).

### Parameter determination

To find optimal parameter values, we fit the loop extrusion model to CTCF ChIA-PET data by fixing two of the three parameters and varying the remaining one. The best-fitting parameter is defined to be the one reaches maximum AUPRC. This method is effective since the nonlinearly in this model makes it hard to perform a maximum likelihood estimation by canonical methods like logistic regression. Taking GM12878 as an example, by fixing dissociation constant <*K_d,i_*>/[CTCF] (*a*) and Cohesin processivity λ, we found *w* value of the best agreement with data is 3. By fixing w and λ, we found the optimal <*K_d,i_*>/[CTCF] is 8.5. Optimal *w* and <*K_d,i_*>/[CTCF] for HeLa is quite similar, 2.8 and 8. For λ, the performance of our model monotonically increases when λ is larger, and asymptotically approaches to the performance of model without this distance-associated exponent term (*D_ij_*=1). We also performed a grid search over these three parameters and found high performance in a broad range around this single optimal set of values.

### CTCF ChIA-PET data processing

GM12878 and HeLa CTCF ChIA-PET data were taken from a published dataset^16^. ChIA-PET2 pipeline with long read mode was used to process data and identify loops^30,69^. One mismatch was allowed in identifying reads with linkers in linker filtering step. Default parameters were used for other steps. Loops are required to be supported by at least 4 PETs for GM12878 and 3 PETs for HeLa. We further constrained CTCF interactions to be within 1 million bp (Mb), as over 96% of loops fell into this range.

### CTCF ChIP-seq data processing

CTCF ChIP-seq of GM12878, HeLa and K562 was obtained from the ENCODE portal. Reads were aligned with BWA to the hg38 reference genome^70^. Peaks were called by MACS2 with default parameters^71^.

### CTCF motif analysis of ChIP-seq data

The position weight matrix of human CTCF was download from JASPER^72^. STORM with default parameters was used to identify the strongest CTCF motif and the corresponding strand for each CTCF binding site, to select the value of the orientation parameter *w*.

### Boosting model

An ensemble-learning-based boosting model was construct with the python Xgboost package. The model consisted of 50 trees, each with maximum depth of 5 layers. The components of the loop extrusion model are used as input features independently. We performed 10-fold cross validation on segregated chromosomes, and averaged performance to account for randomness between chromosomes. Xgboost is able to perform better (Fig 3ab) than the loop extrusion model on the limited subset of features (BI, Ori, Dist). We believe this is because when retrained on this subset, Xgboost is learning an appropriate distance weighting in the absence of loop competition (LC), while for the loop extrusion model we used the optimal *λ* determined using all features. As discussed in Fig 5, loop competition and distance are correlated features, and Xgboost can learn some of the effects of loop competition by regressing on distance.

### Lollipop model

Lollipop is a previously published random forest model which can accurately predict CTCF interaction specificity using 77 features^26^. It has been evaluated on the same CTCF ChIA-PET dataset processed in a very similar method. Therefore, we directly compare the AUROC and AUPRC with Lollipop. Although the original setup of Lollipop training used random test sets, the performance was similar when we reran with chromosomal test sets, with AUPRC=0.88 (random) and 0.86 (chromosomal) for GM12878, and AUPRC=0.90 (random) and 0.89 (chromosomal), so the overfitting due to shared features^59^ is minimal for this training data set.

### Micro-C data processing

A total of 15,945 loops were called from 2.6B reads of mESC Micro dataset^36^. Chromatin loops were identified by using HiCCUPS^6^. Loops were called at 1Kb resolutions at peak size = 4Kb, window size = 10Kb, distance to merge = 2.5Kb and FDR<0.1.

### Distance and loop competition matched sampling of ChIA-PET dataset

The effects of CTCF binding intensity, orientation, distance, and loop competition on CTCF loop formation are quantified separately by four terms *p_i_*p_j_*, *w_ij_*, *D_ij_* and *LC_ij_*. For each positive interaction loop, we define a distance matched non-interacting CTCF pair to be one with the same CTCF motif orientation, with the difference of *p_i_*p_j_* and *D_ij_* between the two loop pairs within a factor of two. Therefore, the difference of distance between them is controlled, while the magnitude of the loop competition term *LC_ij_* is not. Similarly, a loop competition matched non-interacting CTCF pair is one with the same CTCF motif orientation, with the difference of *p_i_*p_j_* and *LC_ij_* between them within a factor of two. These selection procedures generate two positive and negative CTCF pair sets with either matched distance or matched loop competition. We then evaluate our model’s ability to accurately distinguish the positive and negative pairs in both sets, when including either loop competition or distance terms in our model.

### Predicting CRISPR perturbation effect

mESC CTCF ChIP-seq data were taken from GSE72720. The loop extrusion model was built and interacting CTCF pairs are predicted quantitatively, with *K_d_* = 8.5, w = 3, λ = 3,000,000. The effect of CRISPR deletion and inversion of CTCF motif on CTCF binding intensity are taken from^9^. For 4C signal, we calculated the ratio of read counts per kilobase between 20kb bins centered around the perturbed CTCF binding site and 200kb random genomic regions. The change of binding intensity and orientation are then integrated into model to determine the resulting interaction probability.

### Population Hi-C data processing

Normalized Hi-C contact matrices of lymphoblastoid cell lines (LCLs) were taken from^37^. Briefly, Hi-C was performed on LCLs of 20 individuals with previously cataloged genetic variation. Reads were aligned to hg19 reference genome with BWA-MEM as described in^55,70^. Raw counts of contact matrices were normalized to correct for known biases with HiCNorm^73^, as described in^37^.

### WAPL knockout model

WAPL is known as Cohesin unloading factor, as it removes Cohesin from binding with chromatin fiber. It’s been reported that WAPL knockout increase cross-TAD chromatin interaction frequency and extends the size of chromatin loops. We hypothesize that this effect is due to the longer residence time of Cohesin on the chromatin fiber in the context of a WAPL knockout, which allows Cohesin to pass through existing loop boundaries (e.g. CTCF or other Cohesin) with some small probability, *s*, following Ref. ^29^. This pass-through probability attenuates the influence of loop competition and facilitates longer loop formation. The pass-through probability, *s,* thus reduces the loop competition effect of each overlapping CTCF loop by a factor of *1 - s*, and results in a larger loop interaction probability *p_ij_*, which explains the increased loop number under WAPL knockout. The effect of Cohesin passing through is especially strong for distant CTCF pairs, as their interactions are likely to be affected by more competing loops than nearby CTCF pairs. Therefore it also explains the experimentally observed formation of higher order loop interactions (Supp Fig.7d), and consistent with the shift of CTCF loop length distribution to the higher end under large *s* (Supp Fig.7c).

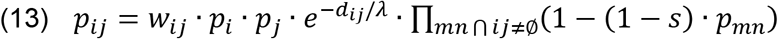

### Cell-type specific CTCF loop identification

Loops from two cell lines are defined to be common if both anchors overlap, if not, we classify them as cell-type specific. We compared the top 10,000 loops in HeLa with all loops in GM12878, and found 956 HeLa-specific loops. Similarly, we compared the top 10,000 loops in GM12878 with all loops in HeLa, and found 2,257 GM12878-specific loops.

## Code Availability

Source code is available for download from https://github.com/wangxi001/Loop-Extrusion-Model or http://doi.org/10.5281/zenodo.4404848.^74^

## Data Availability

Training data available for download from https://github.com/wangxi001/Loop-Extrusion-Model or or http://doi.org/10.5281/zenodo.4404848.^74^

## Acknowledgements

We thank the following members of the Beer Lab for discussion and useful comments on the manuscript: Dustin Shigaki, Jin-Woo Oh, and Milad Razavi-Mohseni. We thank J. Nasser and J. Engreitz for kindly providing a summary table of the enhancer-promoter interactions from references^44–54^. This work was supported by NIH grants HG009380 and HG007348 to MB.

## Author Information

Johns Hopkins University School of Medicine,

733 N. Broadway, Baltimore, MD 21205

Wang Xi & Michael A Beer

Correspondence to.

## Author Contributions

XW and MB designed the study, XW analyzed the data, and XW and MB wrote the paper.

