## Supplemental Figures and Tables for "Loop competition and extrusion model predicts CTCF interaction specificity"

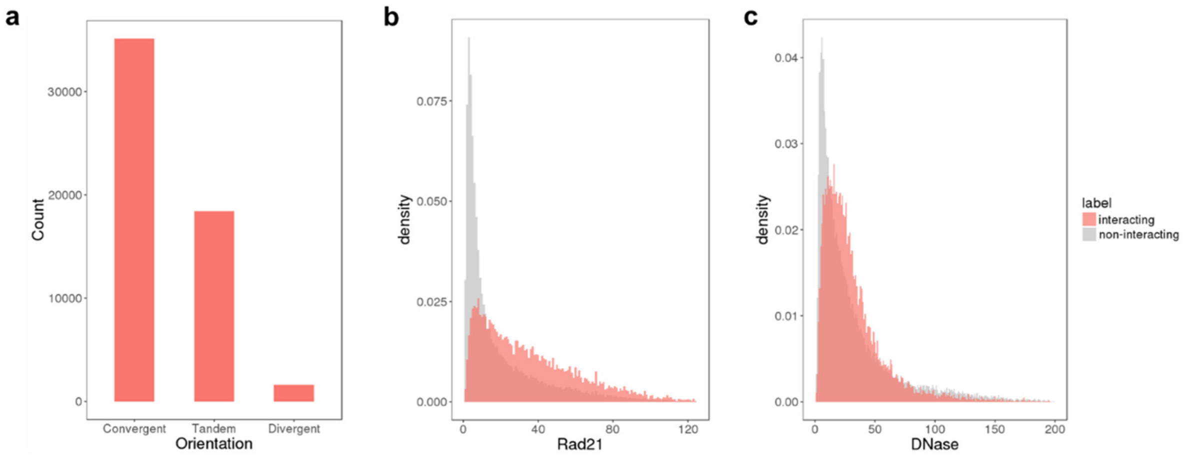


**Supplementary Figure 1. Feature value distributions for interacting and non-interacting CTCF pairs.** (a) Convergent CTCF motifs outnumber tandem or divergent CTCF motifs in interacting CTCF pairs. (b)-(c) Distribution of Rad21 ChIP-seq and DNase-seq signal between interacting and non-interacting CTCF pairs.

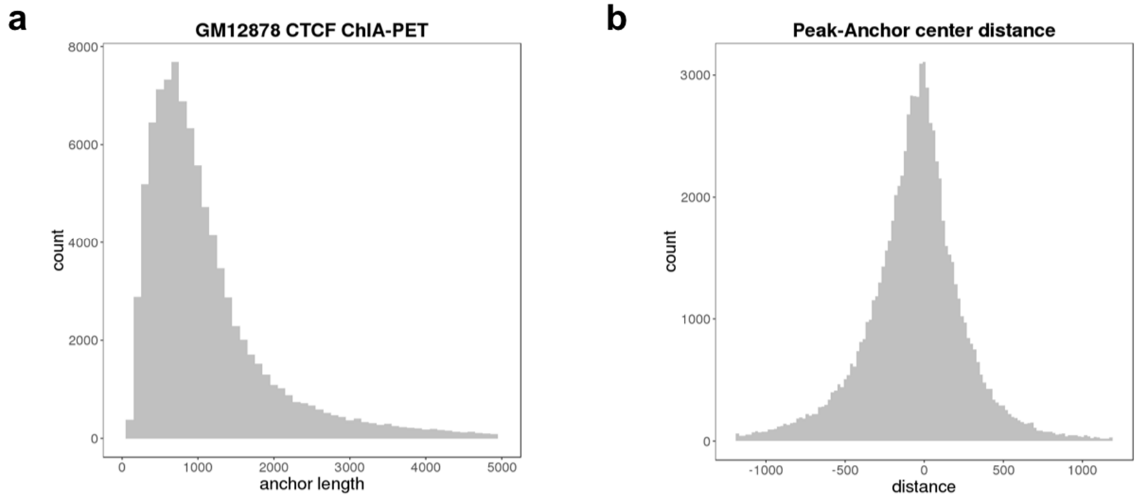


**Supplementary Figure 2. Additional properties of CTCF ChIA-PET data.** (a) Length distribution of GM12878 ChIA-PET loop anchors. (b) CTCF ChIP-seq peaks and ChIA-PET anchors are relatively close, and centered around each other.

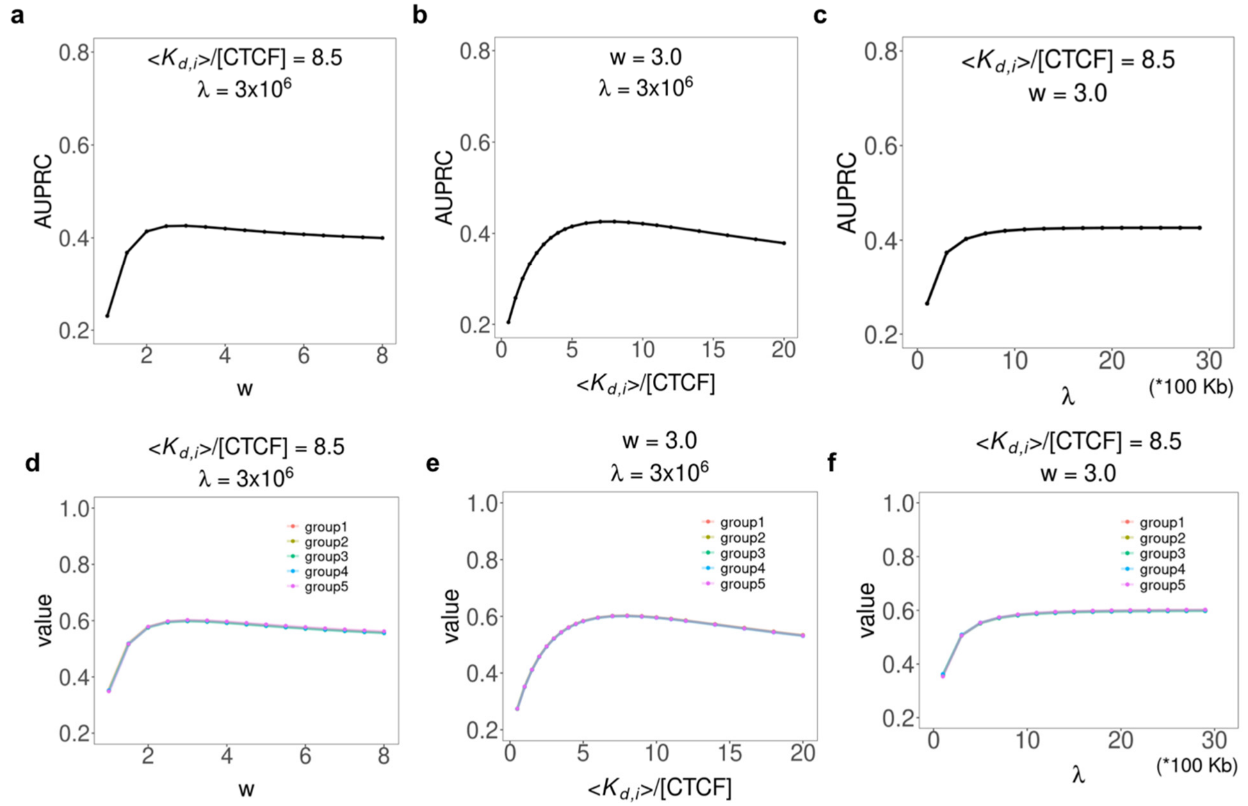


**Supplementary Figure 3. Parameter determination of loop extrusion model on from training on HeLa CTCF ChIA-PET data.** (a)-(c) Parameter search for loop extrusion model. Optimal parameter values are very close to the results on GM12878 ChIA-PET data. (d)-(f) Chromosomal-segregated determination of model parameters on GM12878. Model parameters are determined on each training set after separating all data into five chromosomal cross-validation training and testing sets.

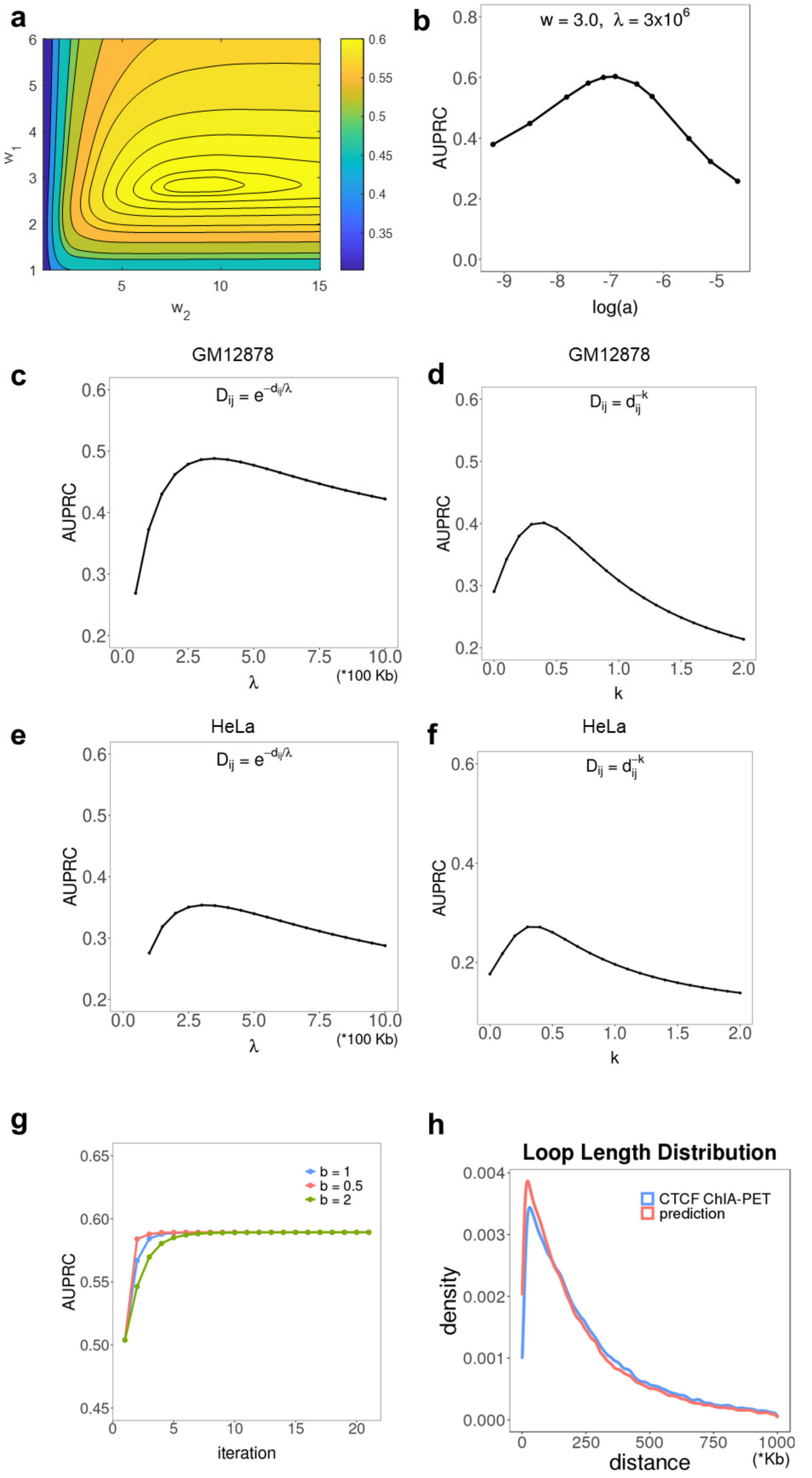


**Supplementary Figure 4. Model performance is robust to specific parameter choices and model assumptions.** (a) Model performance after relaxing $w_{ij}$ for convergent, tandem and divergent loops from $1$, $1/w$, $1/w^2$, into $1$, $1/w_1$, and $1/w_2$. The best-fitting model is achieved at $w_1$

= 3 and $w_2$=9, so we used the simpler model with one parameter, $w$=3. (b) Model performance is robust to the functional form used to map CTCF ChIP-seq signal to binding probability. Using $p_i = \tanh(ax)$, optimal performance at log(a) = -7 is AUPRC=0.6, which is very similar to performance with Eq. (1). (c)-(f) Model performance under distance-dependent exponential decay and power law decay are similar, with slightly improved performance for exponential. (g) Iterative training of model quickly converges to optimal performance under different b, and yields similar performance to Eq. (4). (h) Loop length distribution of ChIA-PET annotated loops and predicted interacting loops. In general, model performance is quite similar in a broad range of parameter choices near their optimal values.

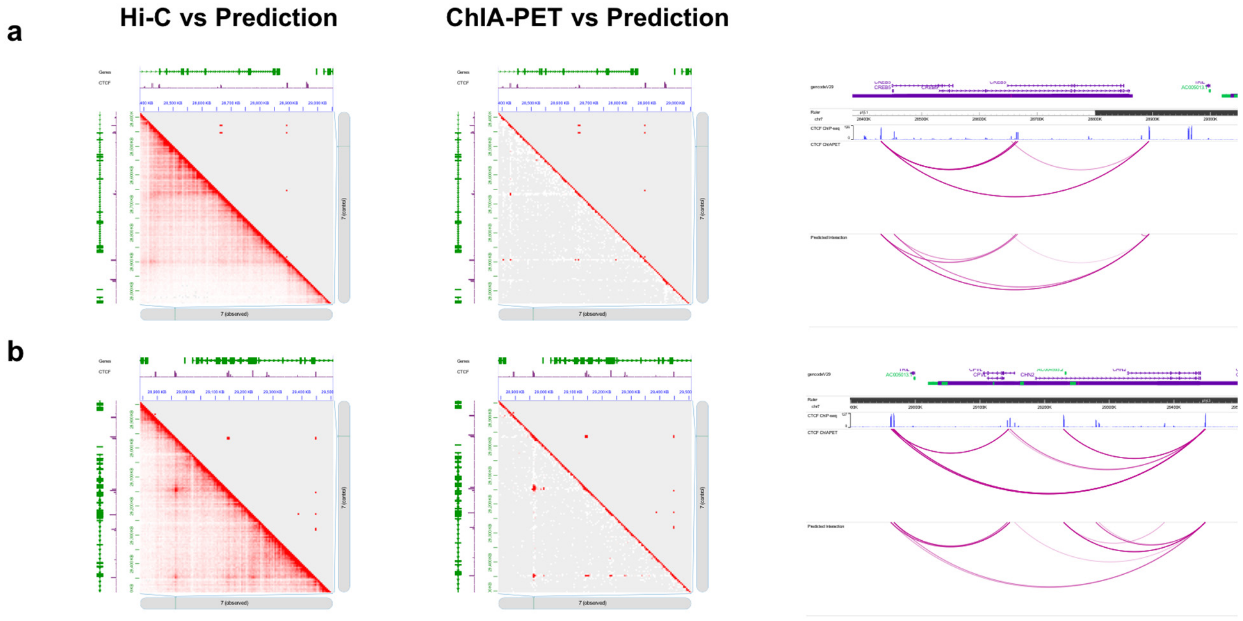


**Supplementary Figure 5. Comparison between Hi-C, ChIA-PET and model prediction.** Loop extrusion model accurately predicts interacting and non-interacting CTCF pairs for two additional loci near (a) CREB5 and (b) CPVL and CHN2, as in Fig. 3.

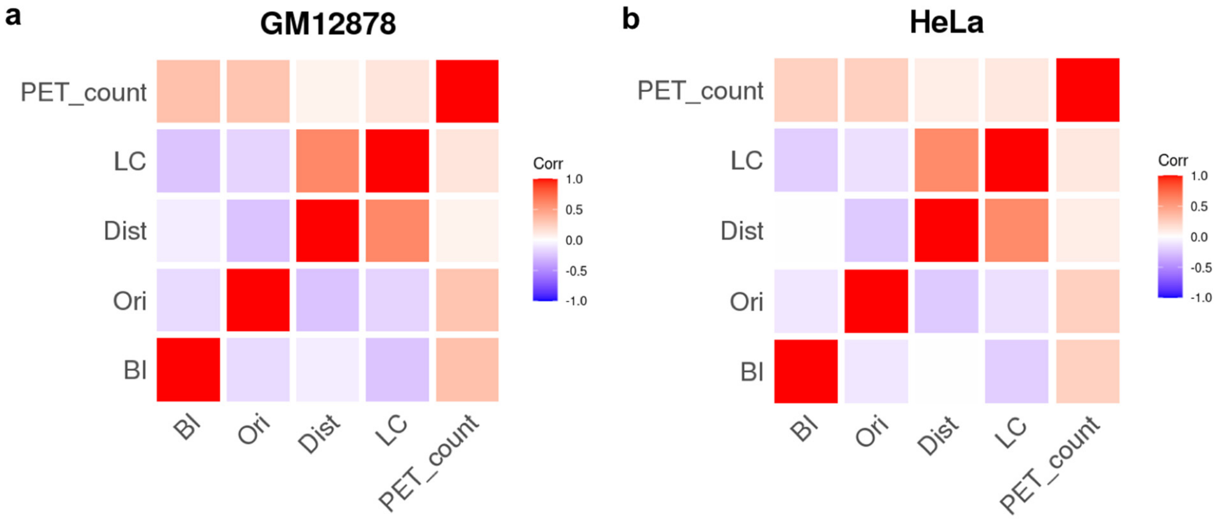


**Supplementary Figure 6. Feature correlation among interacting CTCF pairs.** (a)-(b) Correlation between CTCF binding intensity, CTCF motif orientation, distance, loop competition and PET count (log scale) for interacting CTCF pairs. Distance has the weakest correlation with PET count among the four features.

**a**

Ctrl: $$p_{ij} = w_{ij} \cdot p_i \cdot p_j \cdot e^{-d_{ij}/\lambda} \cdot \prod_{mn \bigcap ij \neq \emptyset} (1 - p_{mn})$$

WAPL KO: $$p_{ij} = w_{ij} \cdot p_i \cdot p_j \cdot e^{-d_{ij}/\lambda} \cdot \prod_{mn \bigcap ij \neq \emptyset} (1 - (1 - s) \cdot p_{mn})$$

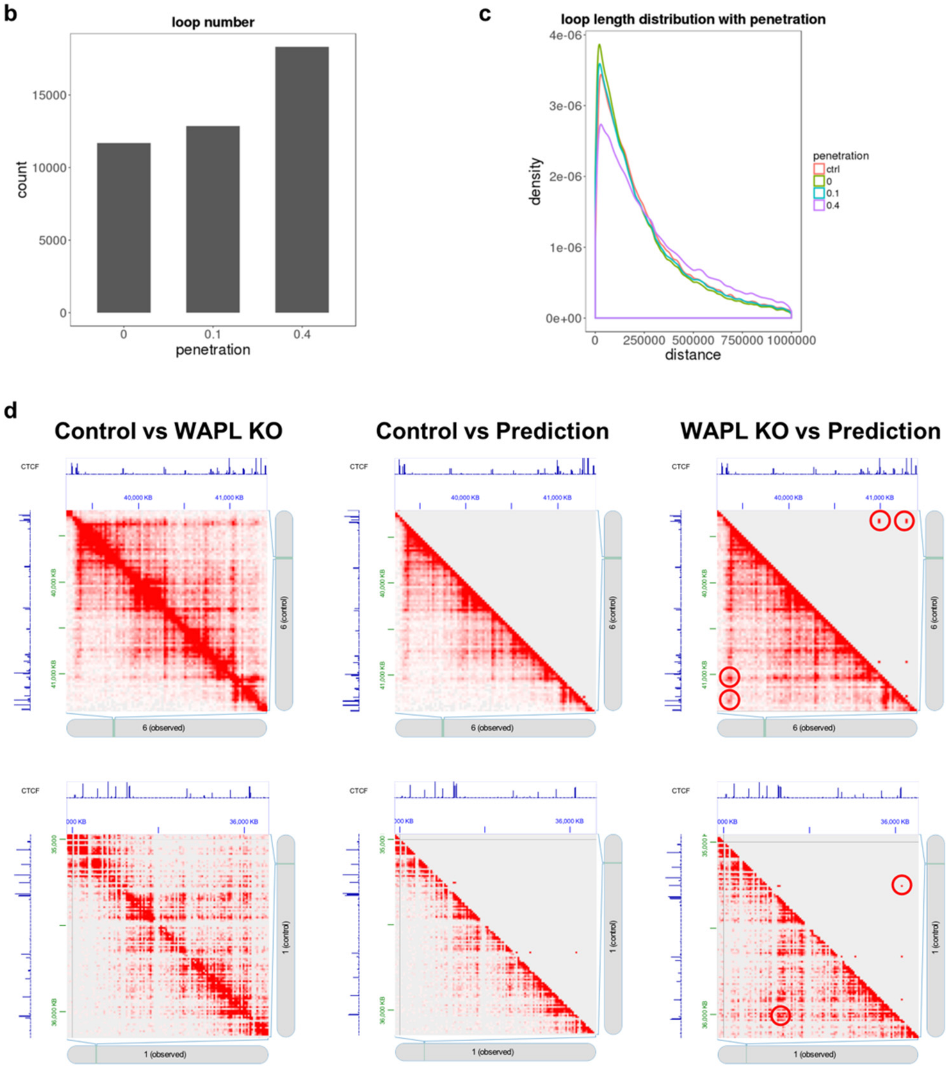


**Supplementary Figure 7. WAPL knockout increases overall CTCF loop length and number. (**a) Loop extrusion mathematical model accounting for WAPL knock out conditions. The parameter $s$ indicates the probability of Cohesin passing through another bound Cohesin. (b) Loop count increases with higher probability of Cohesin pass-though. (c) Longer CTCF loops tend to be formed at higher probability of Cohesin pass-through. (d) Comparison between Hi-C data and model prediction before and after WAPL knockout. Our model successfully predicts formation of long distance CTCF loops.

a

| GM12878 | AUROC (LE model) | AUPRC (LE model) | AUPRC (xgboost) |
|---|---|---|---|
| CTCF | 0.804 | 0.155 | 0.157 |
| Orientation | 0.745 | 0.105 | 0.108 |
| Distance | 0.746 | 0.109 | 0.124 |
| Loop competition | 0.807 | 0.140 | 0.144 |
| CTCF & Orientation | 0.883 | 0.290 | 0.293 |
| CTCF & Orientation Distance | 0.896 | 0.345 | 0.512 |
| CTCF & Orientation Loop competition | 0.953 | 0.601 | 0.602 |
| All | 0.954 | 0.600 | 0.611 |

b

| HeLa | AUROC (LE model) | AUPRC (LE model) | AUPRC (xgboost) |
|---|---|---|---|
| CTCF | 0.759 | 0.088 | 0.088 |
| Orientation | 0.754 | 0.071 | 0.073 |
| Distance | 0.778 | 0.078 | 0.089 |
| Loop competition | 0.816 | 0.097 | 0.100 |
| CTCF & Orientation | 0.857 | 0.177 | 0.176 |
| CTCF & Orientation Distance | 0.873 | 0.221 | 0.369 |
| CTCF & Orientation Loop competition | 0.938 | 0.423 | 0.424 |
| All | 0.940 | 0.426 | 0.435 |

**Supplementary Table 1. Perfromance of loop extrusion model and a less constrained model using the same features (xgboost).**

| Model | AUPRC |
|---|---|
| Baseline model (CTCF, orientation, distance) | 0.512 |
| Baseline model + DNase-seq | 0.530 |
| Baseline model + Rad21 | 0.556 |
| Baseline model + Znf143 | 0.535 |

**Supplementary Table 2. Adding DNase-seq, Rad21, or Znf143 signal strength as features does not improve prediction CTCF interaction significantly.**
